## Supporting Information for "Enhancing extracellular vesicle cargo loading and functional delivery by engineering protein-lipid interactions"

**Contents:**

- Figures S1-S16
- Tables S1-S8

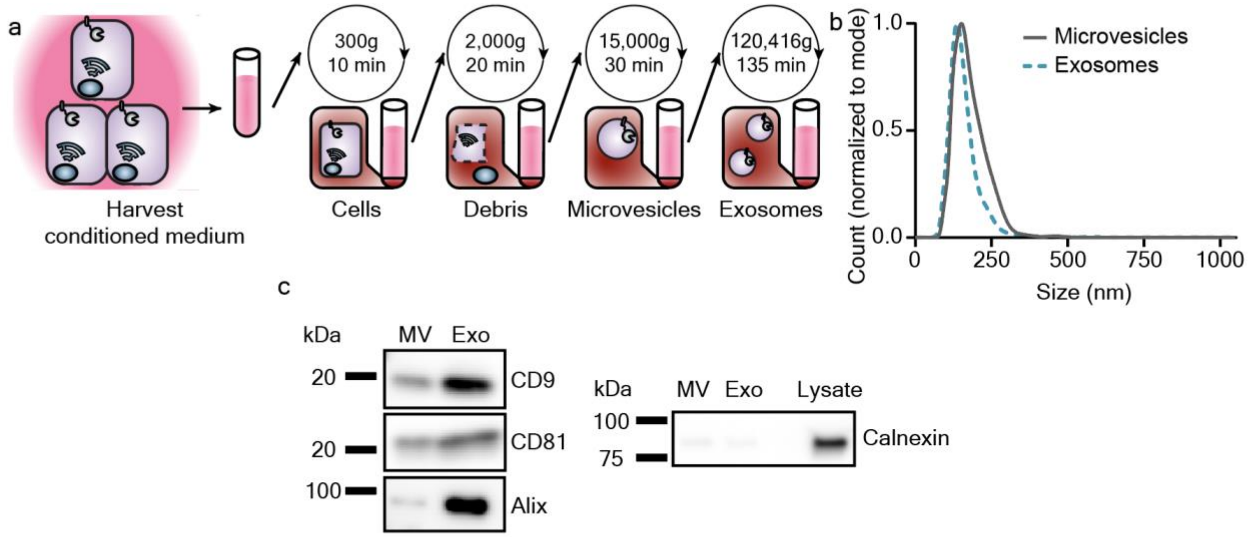


**Supplementary Figure 1. Extracellular vesicle (EV) populations from HEK293FT cells demonstrate classical EV characteristics. a** Cartoon depicting the process for isolating two EV subtypes, microvesicles (MV) and exosomes (Exo), from conditioned cell culture medium. **b** Representative histogram of particle sizes from the EV subpopulations described in **a** as determined by nanoparticle tracking analysis. **c** Western blots on HEK293FT EV samples and cell lysates for EV markers CD9, CD81, and Alix and the endoplasmic reticulum-associated (non EV-associated) marker, Calnexin (n = 1). Equal numbers of EVs were added for each blot. For the Calnexin blot, 4.3 x $10^8$ EVs were used per well, and 2.7 µg of lysate was added.

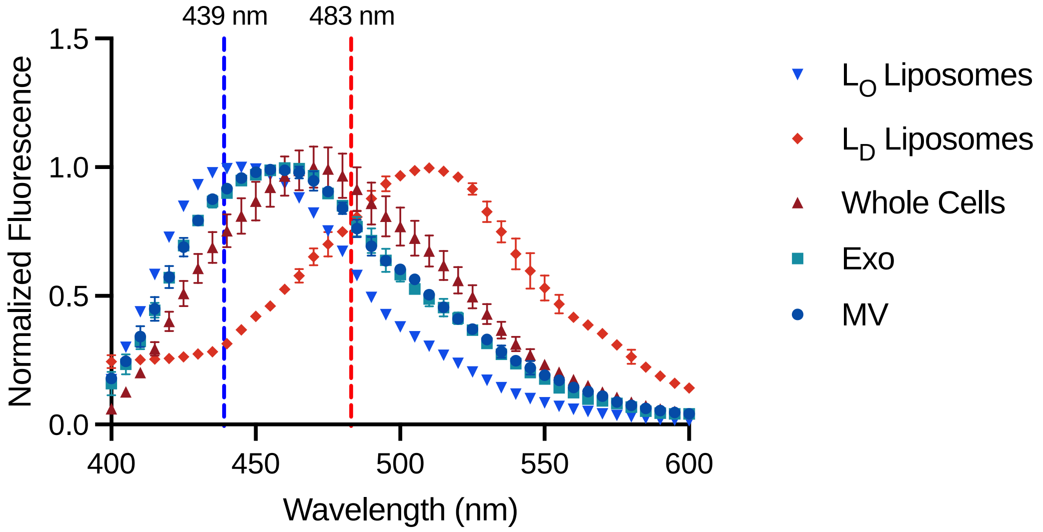


**Supplementary Figure 2. Laurdan spectra of samples in Fig. 1c demonstrate that microvesicles (MV) and exosomes (Exo) are more similar in lipid order to ordered liposomes ($L_O$) than disordered liposomes ($L_D$).** $L_O$ liposomes were composed of 70 mol% DPPC/30 mol% Chol, and $L_D$ liposomes were composed of 70 mol% DOPC/30 mol% Chol. Whole cells refers to HEK293FT cells; the EVs in this experiment were derived from HEK293FTs. Spectra were normalized to the maximum fluorescence. Samples were excited by a 350 nm laser and fluorescence intensities were collected from 400 to 600 nm. Intensities at 439 nm (blue line) and 483 nm (red line) were used to calculate Laurdan generalized polarization (GP) via the formula in the methods section. *n*=3, error bars represent the SEM.

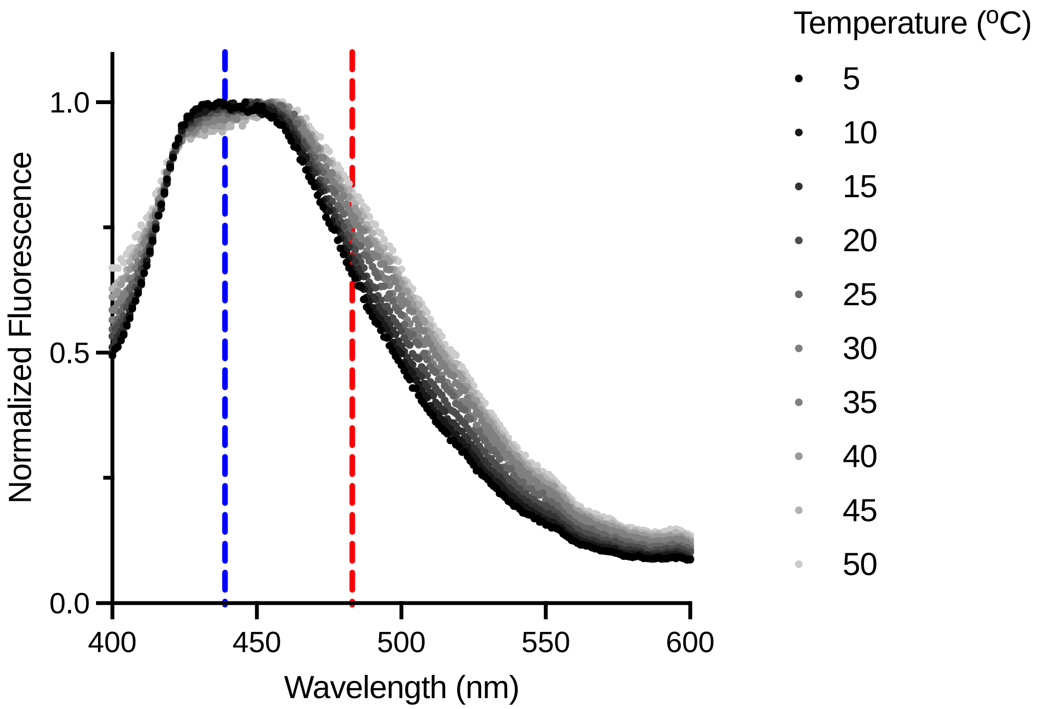


**Supplementary Figure 3. Laurdan spectra of exosomes as a function of temperature demonstrates a characteristic emission profile.** Samples were excited by a 350 nm laser and fluorescence intensities were collected from 400 to 600 nm. Laurdan emission exhibits a slight red shift as temperature increases which reflects the expected decrease in membrane order as temperature increases. Spectra were normalized to the maximum fluorescence. *n*=2, error bars represent the SEM.

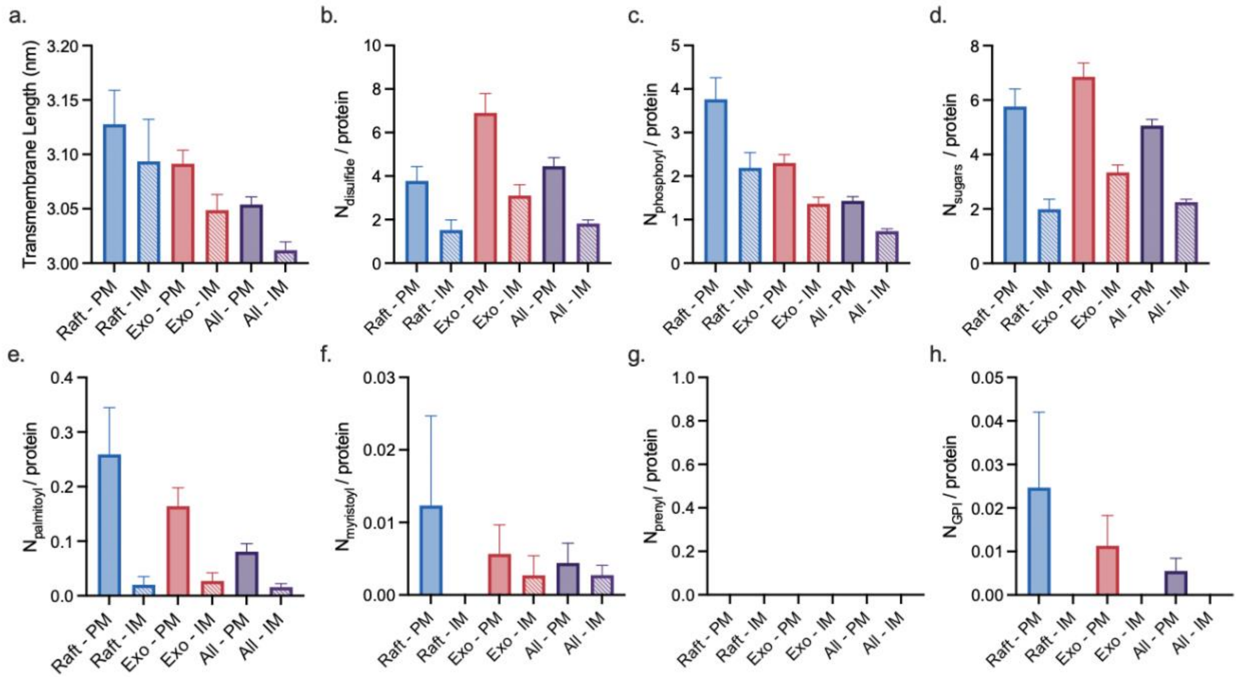


**Supplementary Figure 4. For single transmembrane domain proteins, transmembrane domain length and number of posttranslational modifications can vary between proteins found in lipid rafts (Raft, from Raftprot 2.0), EVs (Exo, from Exocarta), and all human membrane proteins (All, from Swiss-Prot).** Specifically, **a** transmembrane domain length and the average number of **b** disulfides, **c** phosphoryl groups, **d** sugars (glycosylation), **e** palmitoyls, **f** myristoyls, **g** prenyls, and **h** GPI anchors on each protein were calculated. **g** No prenylation of single transmembrane proteins was observed. Proteins were separated by the membrane which they localize to: the plasma membrane (PM) or internal membranes (IM). Error bars represent the SEM.

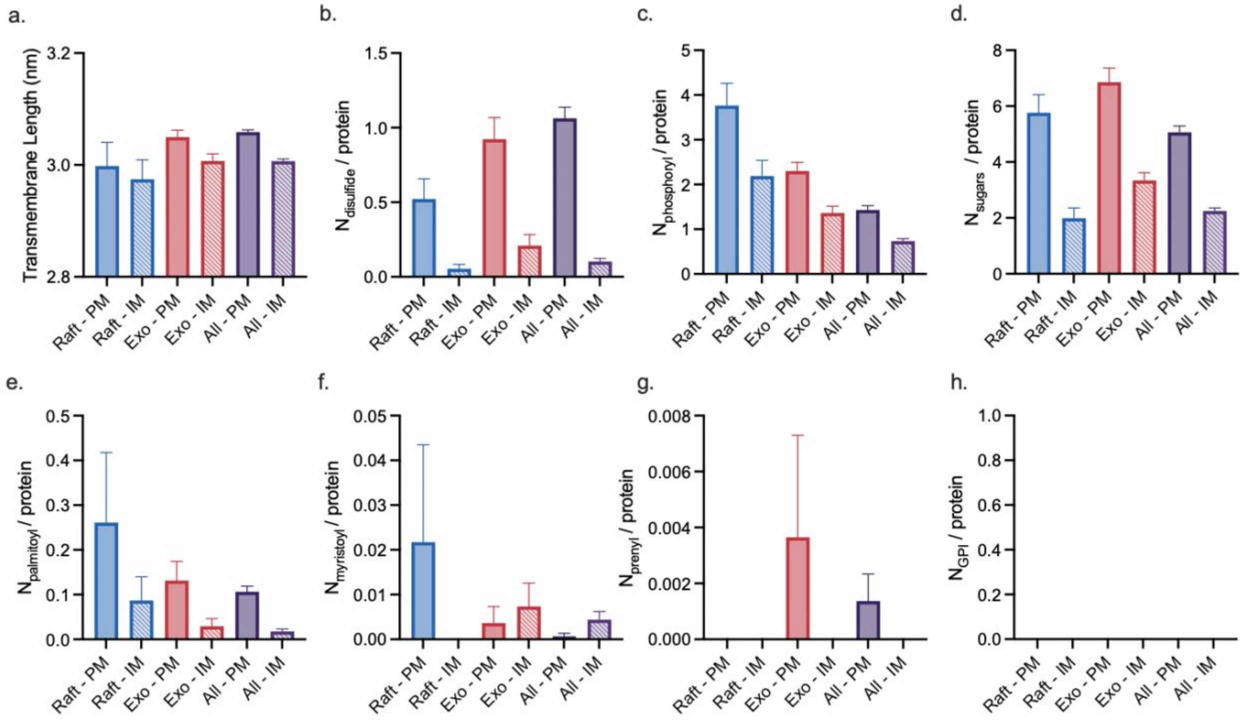


**Supplementary Figure 5. For multi-transmembrane domain proteins, transmembrane domain length and number of posttranslational modifications can vary between proteins found in lipid rafts (Raft, from Raftprot 2.0), EVs (Exo, from Exocarta), and all human membrane proteins (All, from Swiss-Prot).** Specifically, **a** transmembrane domain length and the average number of **b** disulfides, **c** phosphoryl groups, **d** sugars (glycosylation), **e** palmitoyls, **f** myristoyls, **g** prenyls, and **h** GPI anchors on each protein were calculated. **h** No GPI anchors were found on multi-transmembrane proteins. Transmembrane domain length is reported as the average of all transmembrane domains for a single protein. Proteins were separated by the membrane which they localize to: the plasma membrane (PM) or internal membranes (IM). Error bars represent the SEM.

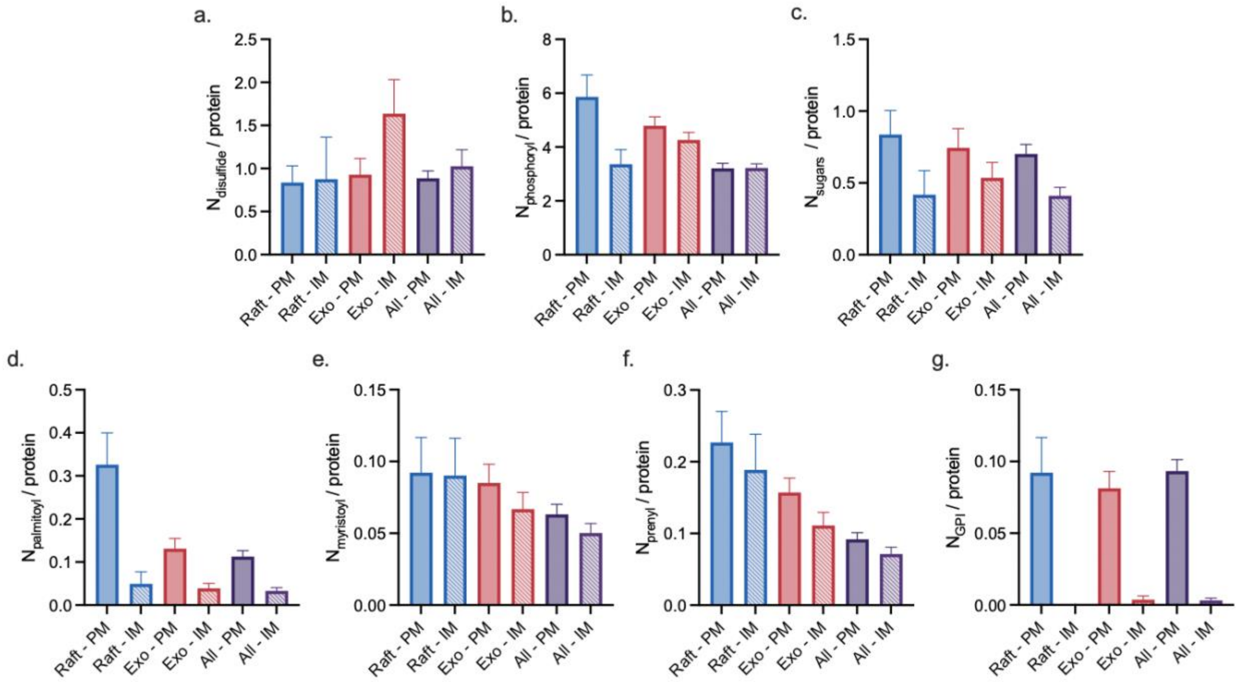


**Supplementary Figure 6. For peripheral membrane proteins, number of posttranslational modifications can vary between proteins found in lipid rafts (Raft, from Raftprot 2.0), EVs (Exo, from Exocarta), and all human membrane proteins (All, from Swiss-Prot).** Specifically, the average number of **a** disulfides, **b** phosphoryl groups, **c** sugars (glycosylation), **d** palmitoyls, **e** myristoyls, **f** prenyls, and **g** GPI anchors on each protein were calculated. Proteins were separated by the membrane which they localize to: the plasma membrane (PM) or internal membranes (IM). Error bars represent the SEM.

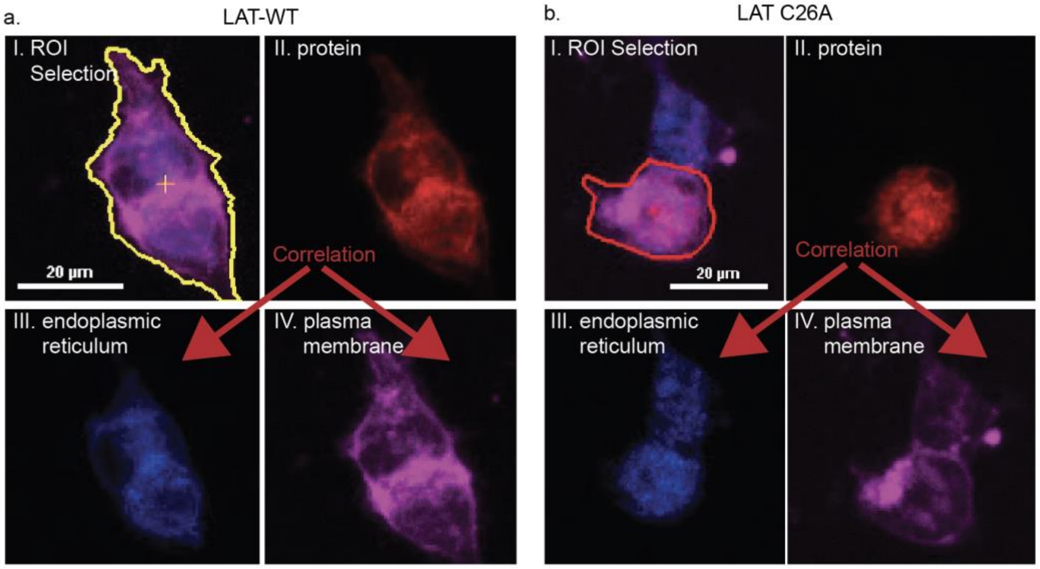


**Supplementary Figure 7. Live-cell protein-localization analysis pipeline used in this study enables quantification of protein colocalization with the plasma membrane or endoplasmic reticulum**. **a, b** Cells transfected with (**a**) LAT-WT and (**b**) C26A LAT are pictured as an example. HEK293FT cells were transfected with each construct and labeled with (III) ER Tracker Blue-White DPX and (IV) Cell Mask Plasma Membrane Dye. Cells were then imaged (I) and colocalization of protein (II) with each dye (endoplasmic reticulum (III) and plasma membrane (IV)) was determined using Nikon Elements Analysis software.

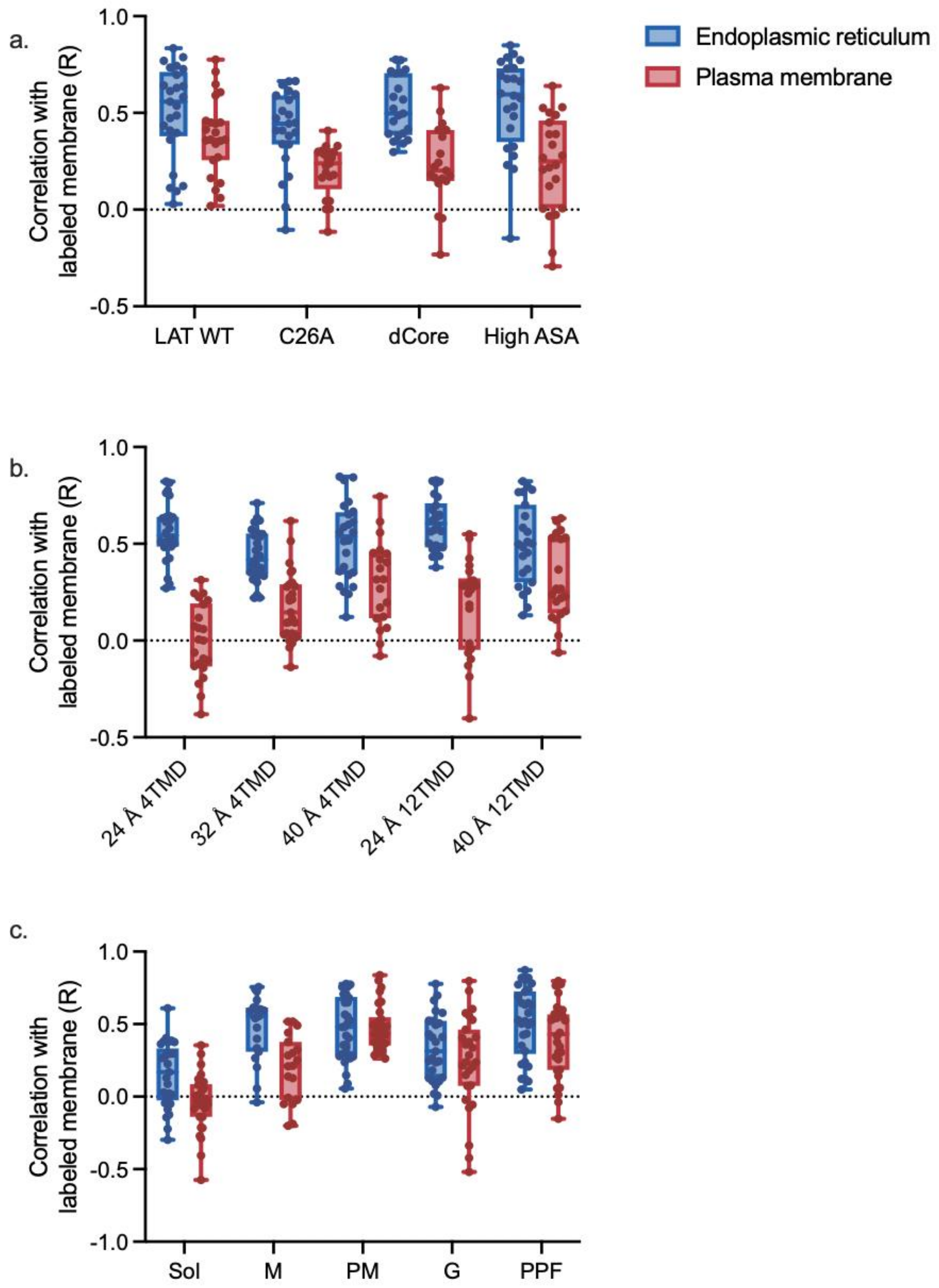


**Supplementary Figure 8. Pearson's coefficients (R) between labeled transgenic protein and the plasma membrane and endoplasmic reticulum enable quantification of protein trafficking. a-c** The plasma membrane was stained with Cell Mask Plasma Membrane Dye, and the endoplasmic reticulum was labeled with ER Tracker Blue-White DPX dyes. The transfected protein construct was either labeled with HaloTag ligand-conjugated dye (TMR) (**a, c**) or directly visualized by mRFP1 fluorescence (**b**). The data in panel **a**, **b**, and **c** correspond to experiments from Fig. 3, Fig. 4, and Fig. 5, respectively. Data are reported in box and whisker plots collected from 30 cells from two independent experiments; each symbol is a single cell.

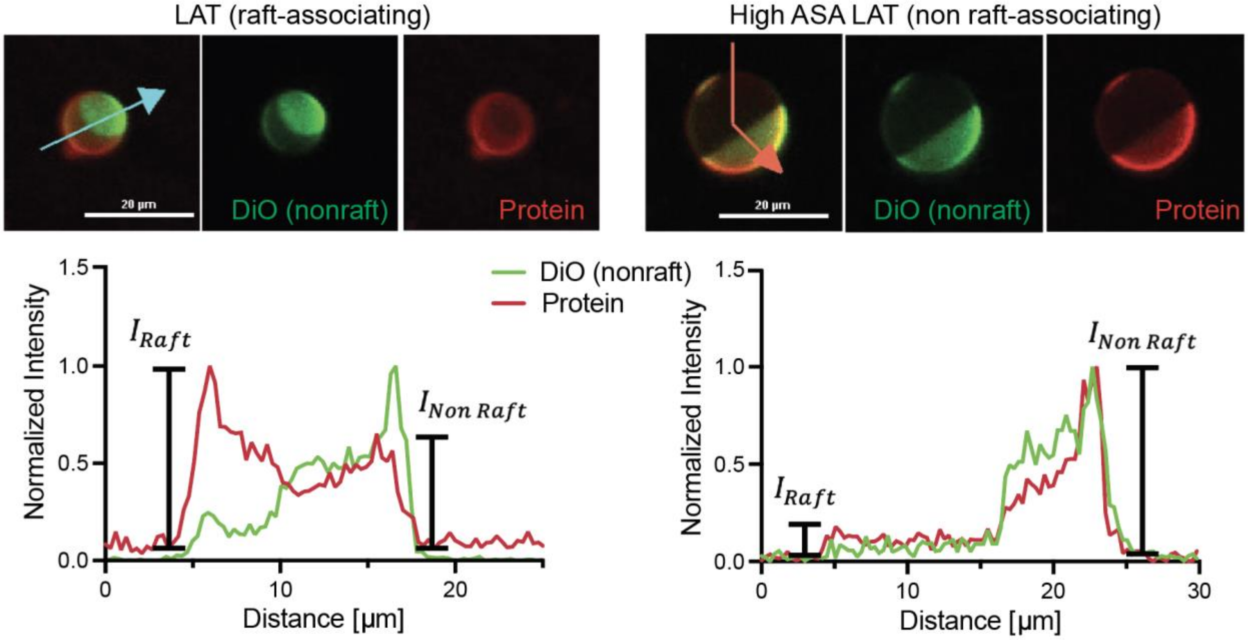


$$L_O\ Enrichment = \frac{I_{Raft} - I_{Non\ Raft}}{I_{Raft} + I_{Non\ Raft}}$$

**Supplementary Figure 9. Giant plasma membrane vesicle (GPMV) analysis pipeline enables an evaluation of protein association with lipid rafts.** HEK293FT cells were transfected with each construct, treated with vesiculation agents, and stained with DiO (non-raft stain). Protein association with lipid rafts was determined by measuring the protein's fluorescence intensity in the raft region (low DiO fluorescence) and nonraft region (high DiO fluorescence) using line scan analysis and using these values to calculate $L_o$ enrichment. $L_o$ enrichment values above 0 indicate proteins prefer lipid rafts.

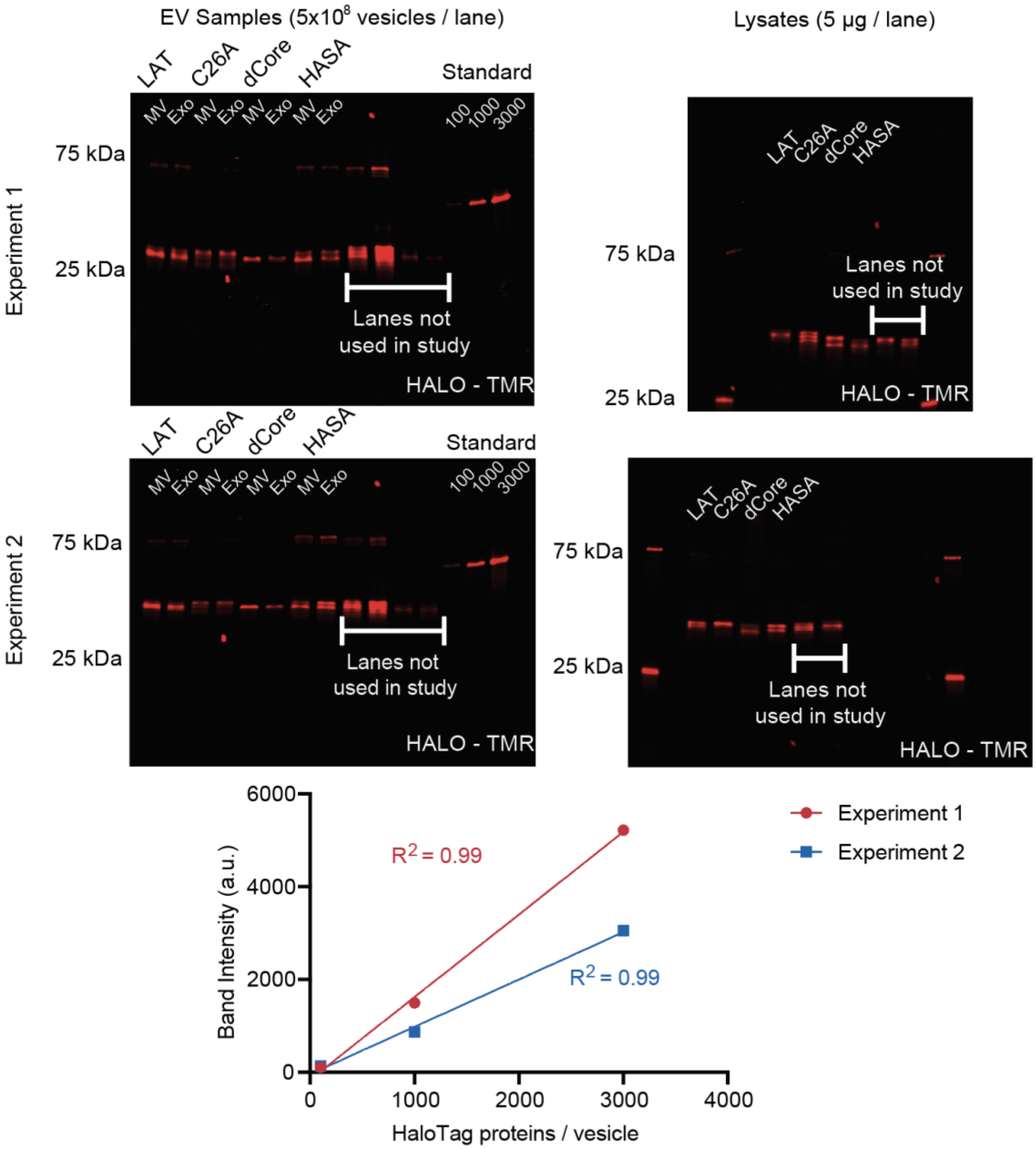


**Supplementary Figure 10. Uncropped protein gels and standard curves generated from LAT transmembrane protein gels show construct expression and loading for cell lysates and EVs, respectively.** Analyzed data is presented in Fig. 3. Values below the standard indicate the number of recombinant proteins added to each well divided by 5 x $10^8$ (the number of vesicles added to the other wells).

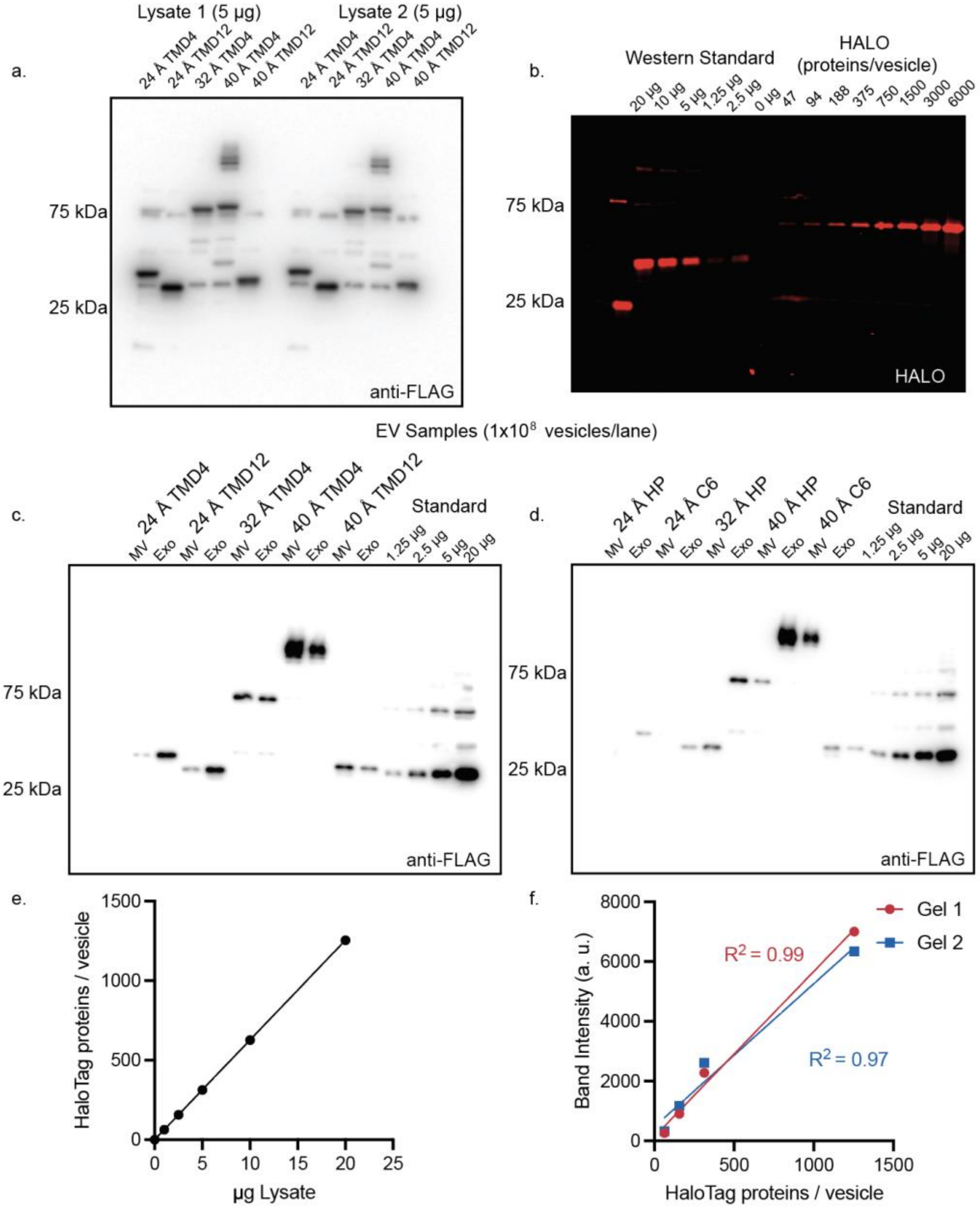


**Supplementary Figure 11. Uncropped western blots and standard curves generated from *de novo* designed transmembrane protein gels show construct expression and loading for cell lysates and EVs, respectively. a** Uncropped western blot of cell lysates. **b** A protein standard with a 3x FLAG tag and HaloTag was run against purified HaloTag (Promega) on an SDS-PAGE gel to quantify the concentration (and equivalent proteins/vesicle) of the protein standard. **c, d** Proteins loaded into vesicles and protein standards were evaluated via western blot. Constructs labeled with 3x FLAG tag. **e** Protein standard and **f** standard curves used to quantify vesicle loading. Analyzed data are presented in Fig. 4d-e.

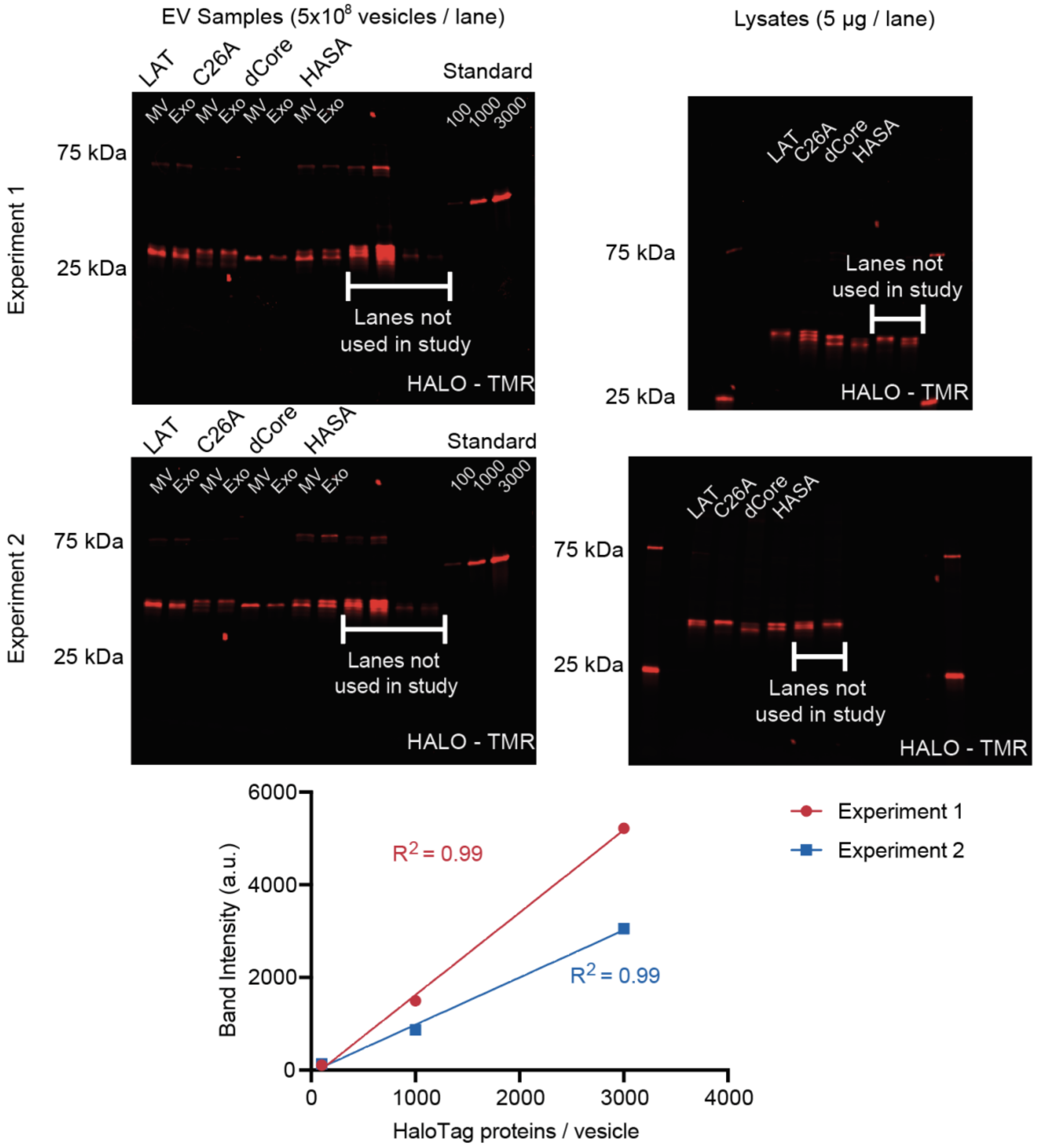


**Supplementary Figure 12. Uncropped protein gels and standard curves generated from peripheral membrane proteins show construct expression and loading profiles for cell lysate and EVs, respectively.** Analyzed data are presented in Fig. 5.

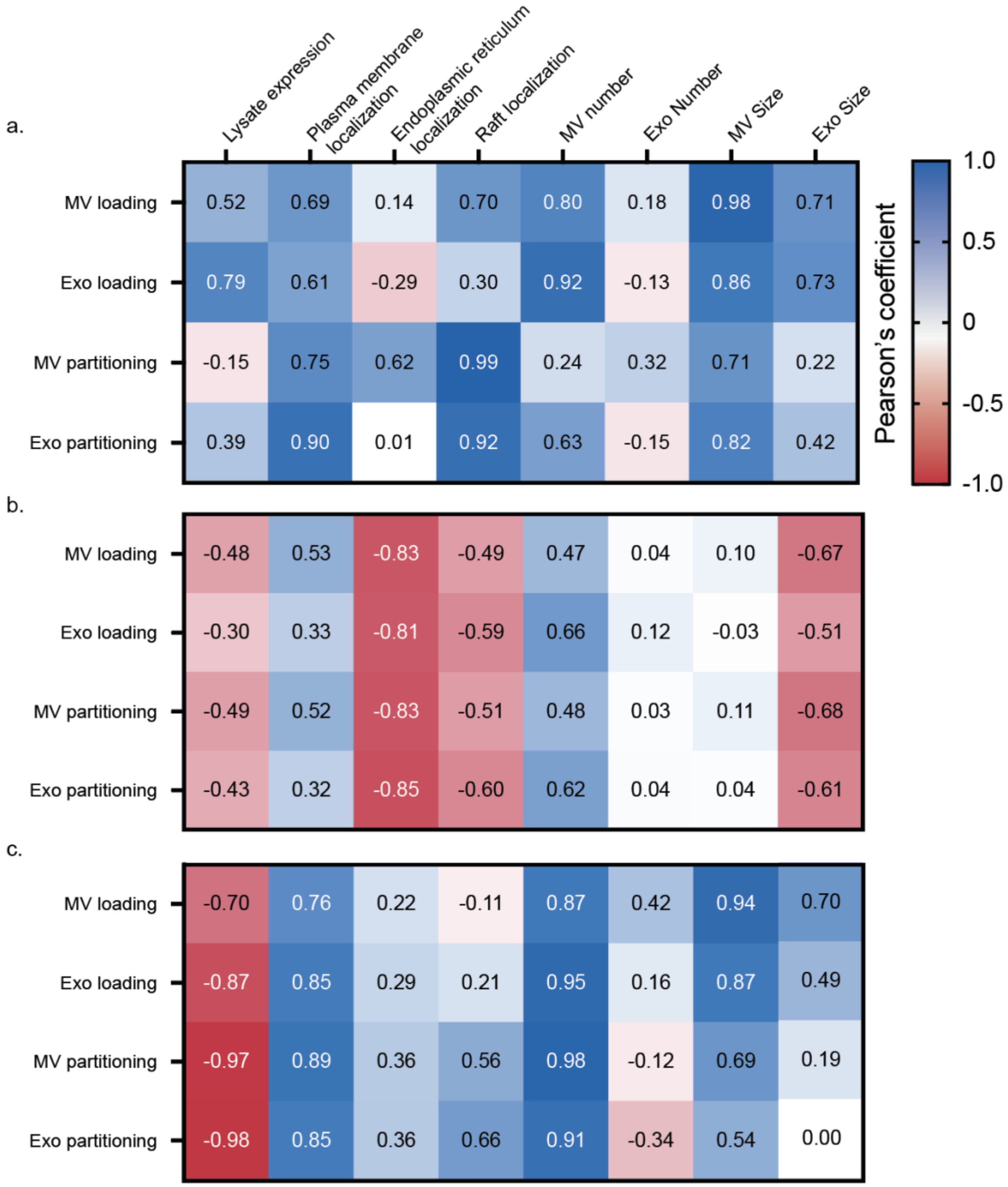


**Supplementary Figure 13. Correlation of protein loading and partitioning into MVs and Exos reveal EV-loading design principles. a-c** All panels quantify correlation using a Pearson's coefficient. **a** Protein loading and partitioning for LAT proteins (LAT WT, LAT C26A, LAT dCore, LAT High ASA). **b** *de novo* designed proteins (24 Å TMD4, 32 Å TMD4, 40 Å TMD4, 24 Å TMD12, 40 Å TMD12). **c** and lipid tagged HaloTag proteins (Sol, M, PM, G, PPF) were correlated with protein expression in lysate, localization to the plasma membrane and endoplasmic reticulum, association with lipid rafts, and vesicle number and size.

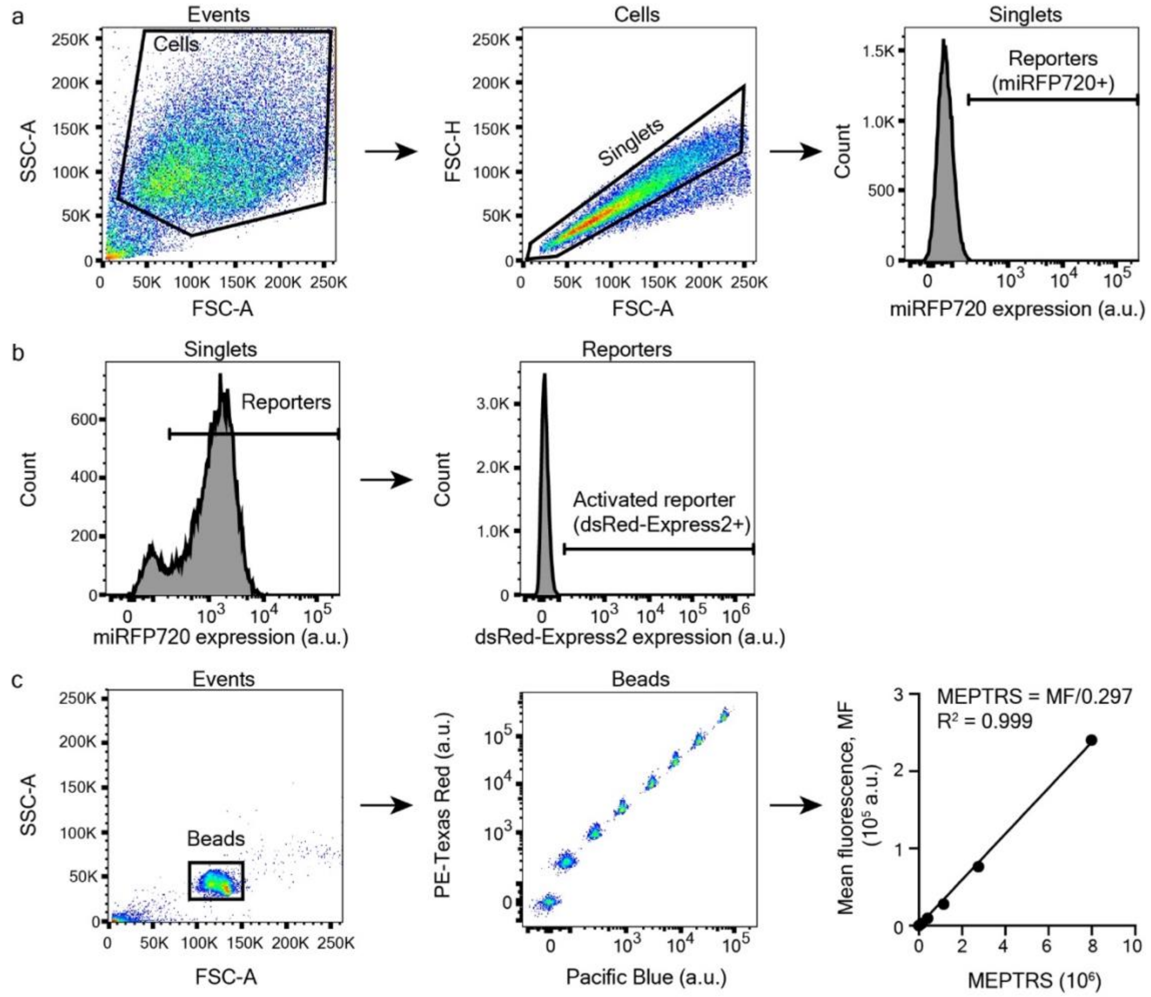


**Supplementary Figure 14. Flow cytometry gating and workflow enable quantification of relevant data (e.g. Fig. 7). a** Events collected during a typical flow cytometry experiment were first gated based on side scatter area (SSC-A) and forward scatter area (FSC-A) to identify HEK293FT cells. Single cells were then identified by forward scatter height (FSC-H) vs FSC-A discrimination. Single color compensation controls were not gated further prior to generating a compensation matrix. For synTF experiments, HEK293FTs expressing no fluorescent proteins were then used to define the miRFP720+ gate (i.e., reporter cells with an active locus), typically such that < 0.1% of HEK293FTs were considered miRFP720+. **b** Left, a sample histogram of single, reporter cells as gated in **a**. Right, reporter cells that did not receive any synTF were used to establish a dsRed-Express2+ gate. **c** To calibrate fluorescence intensity data, UltraRainbow Calibration Particles were run alongside cell samples for each synTF experiment. Beads were first gated based on SSC-A vs FSC-A (left), and then each bead population was gated using two fluorescent channels, typically PE-Texas Red and Pacific Blue. The mean fluorescence (MF) of each grouping was plotted against the vendor-supplied number of equivalent fluorophores (i.e., molecules of equivalent phycoerythrin-texas red, MEPTRs) and a linear regression was performed with the intercept set to 0. A sample regression equation and goodness of fit are provided.

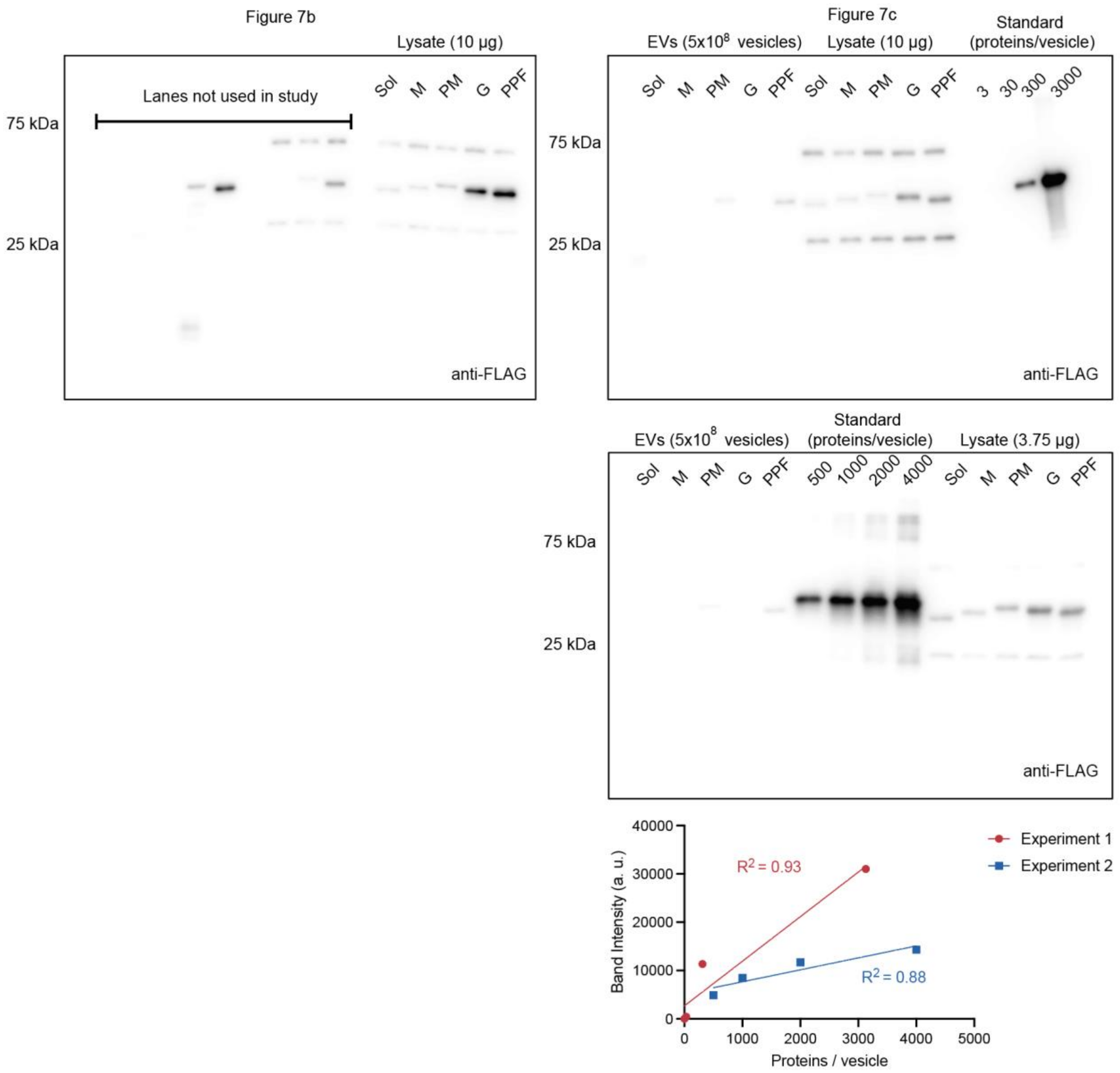


**Supplementary Figure 15. Uncropped western blots and standard curves for EV-mediated synTF delivery experiments show construct expression and loading profiles for lysates from EV-producer cells and EVs, respectively.** Western blots are probing 1x FLAG tag on the constructs. Protein standard is purified p53 with a 1x FLAG tag (R&D Systems). Analyzed data are presented in Fig. 7b (left) and 7c (right).

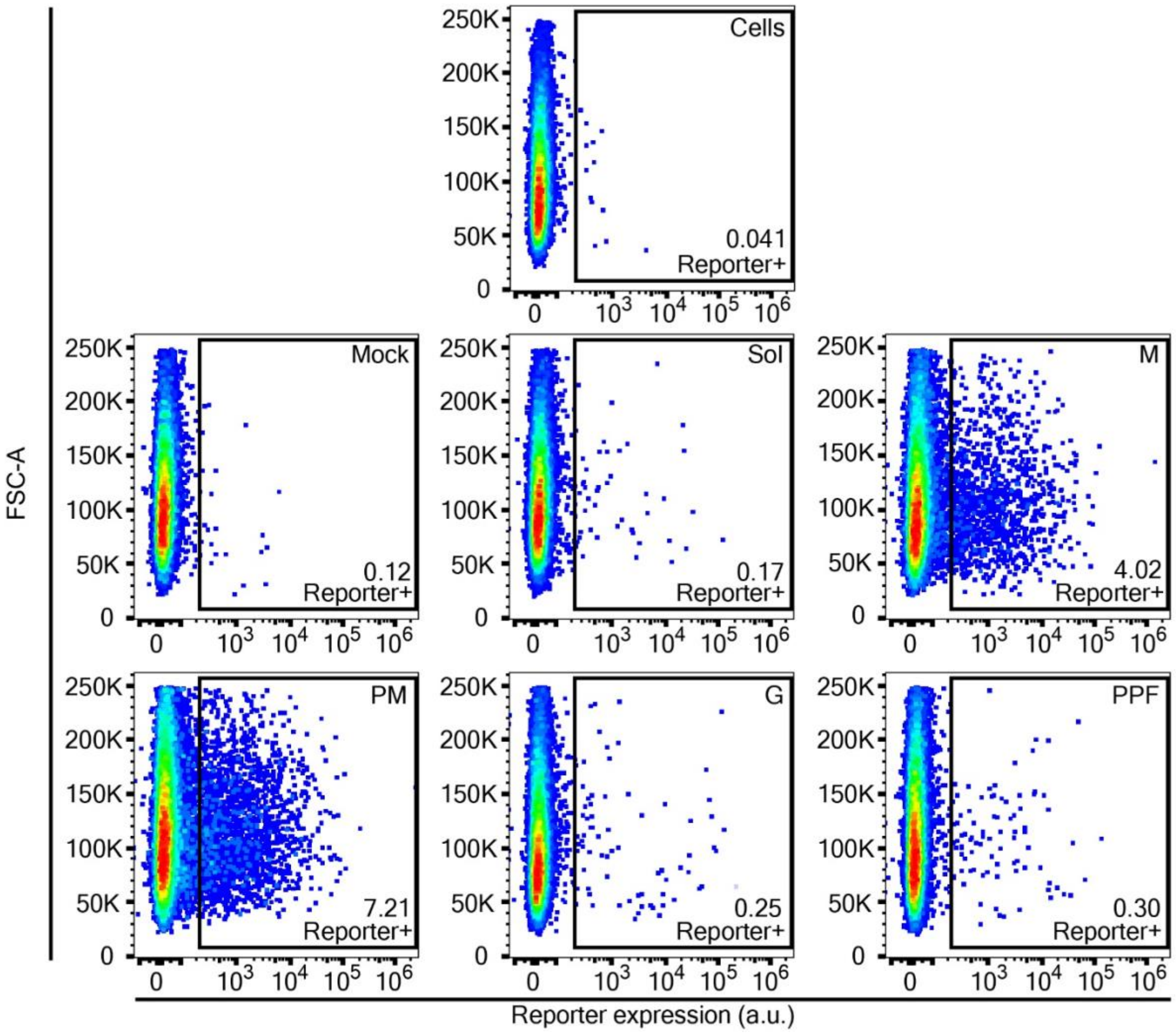


**Supplementary Figure 16. Specific lipidated synTF variants induce reporter expression when delivered via EVs.** Representative dot plots for each condition from the experiment described in Fig. 7. The plots depict forward scatter area (FSC-A) vs reporter expression (dsRed-Express2) for cells treated with EVs containing lipidated synTFs, where each dot represents an individual reporter cell. The lipidated synTF tag designation is denoted in the upper right-hand corner of each dot plot, and the percent of reporter cells activated is denoted in the lower right hand corner of each dot plot. The Mock and PM conditions are also shown in Fig. 7e.

**Supplementary Table 1. Number of proteins analyzed in bioinformatic study for each protein category.**

| | *Protein* | *Single-pass transmembrane protein* | *Multi-pass transmembrane protein* | *Peripheral membrane protein* |
|---|---|---|---|---|
| ***Raft*** | Plasma membrane | 81 | 46 | 122 |
| | Internal membrane | 149 | 82 | 141 |
| ***EV*** | Plasma membrane | 353 | 274 | 541 |
| | Internal membrane | 370 | 272 | 540 |
| ***All Human Proteins*** | Plasma membrane | 904 | 1463 | 1329 |
| | Internal membrane | 1463 | 1360 | 1235 |

**Supplementary Table 2. Number of EVs produced from HEK293FTs transfected with the below constructs, as determined by nanoparticle tracking analysis*.**

| *Construct* | *MVs produced* | *MV Std* | *Exos produced* | *Exo Std* |
|---|---|---|---|---|
| ***Sol HaloTag*** | 8.26E+09 | 1.28E+09 | 1.02E+10 | 1.56E+09 |
| ***M HaloTag*** | 9.32E+09 | 1.41E+09 | 1.08E+10 | 1.66E+09 |
| ***PM HaloTag*** | 1.67E+10 | 2.52E+09 | 1.21E+10 | 1.86E+09 |
| ***G HaloTag*** | 1.40E+10 | 2.14E+09 | 1.34E+10 | 2.10E+09 |
| ***PPF HaloTag*** | 1.53E+10 | 2.34E+09 | 7.04E+09 | 1.24E+09 |
| ***LAT WT*** | 6.97E+09 | 1.12E+09 | 8.16E+09 | 1.27E+09 |
| ***LAT C26A*** | 5.98E+09 | 9.43E+08 | 6.90E+09 | 1.10E+09 |
| ***LAT dCore*** | 5.87E+09 | 8.91E+08 | 8.74E+09 | 1.37E+09 |
| ***LAT High ASA*** | 8.01E+09 | 1.22E+09 | 8.02E+09 | 1.26E+09 |
| ***24 Å 4TM*** | 5.91E+09 | 8.96E+08 | 1.04E+10 | 1.60E+09 |
| ***32 Å 4TM*** | 7.06E+09 | 1.07E+09 | 1.14E+10 | 1.76E+09 |
| ***40 Å 4TM*** | 6.95E+09 | 1.07E+09 | 1.23E+10 | 1.85E+09 |
| ***24 Å 12TM*** | 7.09E+09 | 1.10E+09 | 8.37E+09 | 1.28E+09 |
| ***40 Å 12TM*** | 5.54E+09 | 8.54E+08 | 1.24E+10 | 1.89E+09 |

* n=2

**Supplementary Table 3. EV size for vesicles harvested from HEK293FTs that were transfected with the below constructs, as determined by nanoparticle tracking analysis*.**

| | *MV Size (nm)* | *MV Std (nm)* | *Exo Size (nm)* | *Exo Std (nm)* |
|---|---|---|---|---|
| ***Sol HaloTag*** | 197.4 | 72.9 | 160.1 | 45.9 |
| ***M HaloTag*** | 202.7 | 77.9 | 165.7 | 47.5 |
| ***PM HaloTag*** | 205.7 | 71.3 | 164.5 | 51.6 |
| ***G HaloTag*** | 207.6 | 70.2 | 171.3 | 50.7 |
| ***PPF HaloTag*** | 203.5 | 71.4 | 161.9 | 57.0 |
| ***LAT WT*** | 191.0 | 59.6 | 160.3 | 50.5 |
| ***LAT C26A*** | 184.6 | 68.0 | 158.6 | 51.2 |
| ***LAT dCore*** | 185.8 | 71.6 | 159.6 | 50.1 |
| ***LAT High ASA*** | 190.3 | 72.8 | 161.8 | 49.9 |
| ***24 Å 4TM*** | 173.8 | 64.3 | 151.3 | 51.5 |
| ***32 Å 4TM*** | 174.8 | 64.5 | 148.3 | 46.3 |
| ***40 Å 4TM*** | 172.4 | 57.3 | 155.2 | 46.8 |
| ***24 Å 12TM*** | 181.6 | 65.1 | 148.9 | 51.4 |
| ***40 Å 12TM*** | 173.0 | 61.2 | 153.9 | 45.7 |

*n=2

**Supplementary Table 4. Protein source and sequences for proteins used in this study.**

| TAG | PROTEIN SOURCE | PROTEIN SEQUENCE |
|---|---|---|
| **LAT WT** | LAT | MEEAILVPCVLGLLLLPILAMLMALCVHCHRLP - [POI] |
| **LAT C26A** | LAT-modified | MEEAILVPCVLGLLLLPILAMLMALAVHCHRLP - [POI] |
| **LAT DCORE** | LAT-modified | MEEAILVPCVLGLAMLMALCVHCHRLP - [POI] |
| **LAT HIGH ASA** | LAT-modified | MEEAILVLLLLLLLLLPLLLLLLLLCVHCHRLP - [POI] |
| **24 Å 4TMD** | *de novo* designed | MGSTRTEIIRELERSLREQRVLAIFLLALLIVLLWLLQQLKELLRELERLQREGSSDEDVRELLREIKELVENIVYLVIIIMVLVLVIIALARTQKYLVEELKRQD - [POI] |
| **32 Å 4TMD** | *de novo* designed | MGSTRTEIITRLSFSLLLQLVLAIFLLALLIVLLWLLQQLKELLRELERLQREGSSDEDVRELLREIKELVENIVYLVIIIMVLVLVIIALAVLQMYLVRELKRQD - [POI] |
| **40 Å 4TMD** | *de novo* designed | MGSTRTEIITRLSFSLLLQLVLAIFLLALLIVLLVLLIYLKELLRELERLQREGSSDEDVRELLREIKWLVIVIVALVIIIMVLVLVIIALAVLQMYLVRELKRQD - [POI] |
| **24 Å 12TMD** | *de novo* designed | MTENEIRKLRKLLRIAMFLLVFLLIWTWISLETSKTDDDPSAQSEALVAMSLMLIAASLLIIAKSKLMKSRNG - [POI] |
| **40 Å 12TMD** | *de novo* designed | MTKKIIMVLILLLIIAMLLLVFLLIATVVSLWWSWTDDDPSAISEALVAMSLMLIAASLLIIAISKLLKSKNG - [POI] |
| **M** | Src | MGSSKSKPKDPSQRRNNNNGPVAT - [POI] |
| **PM** | Lyn | MGCIKSKRKDKDLELKLRILQSTVPRARDPPVAT - [POI] |
| **G** | K-Ras | [POI]-FRSDGKKKKKKSKTKCQLL |
| **PPF** | Paralemmin-1 | [POI]-CKCCSIM |

### Supplementary Table 5. DNA sequences for protein loading studies.

| Construct | DNA Sequence |
| --- | --- |
| **WT LAT HaloTag*** | ATGGAGGAGGCCATCCTGGTCCCCTGCGTGCTGGGGCTCCTGCTGCTGCCCATCCTGGCCATGTTGAT<br>GGCACTGTGTGTGCACTGCCACAGACTGCCAGGCTCCGGATCCGGCGGCTCCGAAATCGGTACTGGCT<br>TTCCATTCGACCCCCATTATGTGGAAGTCCTGGGCGAGCGCATGCACTACGTCGATGTTGGTCCGCGCG<br>ATGGCACCCCTGTGCTGTTCCTGCACGGTAACCCGACCTCCTCCTACGTGTGGCGCAACATCATCCCGC<br>ATGTTGCACCGACCCATCGCTGCATTGCTCCAGACCTGATCGGTATGGGCAAATCCGACAAACCAGACC<br>TGGGTTATTTCTTCGACGACCACGTCCGCTTCATGGATGCCTTCATCGAAGCCCTGGGTCTGGAAGAGG<br>TCGTCCTGGTCATTCACGACTGGGGCTCCGCTCTGGGTTTCCACTGGGCCAAGCGCAATCCAGAGCGC<br>GTCAAAGGTATTGCATTTATGGAGTTCATCCGCCCTATCCCGACCTGGGACGAATGGCCAGAATTTGCCC<br>GCGAGACCTTCCAGGCCTTCCGCACCACCGACGTCGGCCGCAAGCTGATCATCGATCAGAACGTTTTTA<br>TCGAGGGTACGCTGCCGATGGGTGTCGTCCGCCCGCTGACTGAAGTCGAGATGGACCATTACCGCGAG<br>CCGTTCCTGAATCCTGTTGACCGCGAGCCACTGTGGCGCTTCCCAAACGAGCTGCCAATCGCCGGTGAG<br>CCAGCGAACATCGTCGCGCTGGTCGAAGAATACATGGACTGGCTGCACCAGTCCCCTGTCCCGAAGCT<br>GCTGTTCTGGGGCACCCCAGGCGTTCTGATCCCACCGGCCGAAGCCGCTCGCCTGGCCAAAAGCCTGC<br>CTAACTGCAAGGCTGTGGACATCGGCCCGGGTCTGAATCTGCTGCAAGAAGACAACCCGGACCTGATCG<br>GCAGCGAGATCGCGCGCTGGCTGTCTACTCTGGAGATTTCCGGCTCCGAATTCGATTACAAGGACCACG<br>ATGGCGACTATAAGGATCACGACATCGACTACAAGGACGATGACGACAAGTGA |
| **LAT C26A HaloTag** | ATGGAGGAGGCCATCCTGGTCCCCTGCGTGCTGGGGCTCCTGCTGCTGCCCATCCTGGCCATGTTGAT<br>GGCACTGgccGTGCACTGCCACAGACTGCCAGGATCCGGCGGCTCCGAAATCGGTACTGGCTTTCCATT<br>CGACCCCCATTATGTGGAAGTCCTGGGCGAGCGCATGCACTACGTCGATGTTGGTCCGCGCGATGGCA<br>CCCCTGTGCTGTTCCTGCACGGTAACCCGACCTCCTCCTACGTGTGGCGCAACATCATCCCGCATGTTG<br>CACCGACCCATCGCTGCATTGCTCCAGACCTGATCGGTATGGGCAAATCCGACAAACCAGACCTGGGTT<br>ATTTCTTCGACGACCACGTCCGCTTCATGGATGCCTTCATCGAAGCCCTGGGTCTGGAAGAGGTCGTCC<br>TGGTCATTCACGACTGGGGCTCCGCTCTGGGTTTCCACTGGGCCAAGCGCAATCCAGAGCGCGTCAAA<br>GGTATTGCATTTATGGAGTTCATCCGCCCTATCCCGACCTGGGACGAATGGCCAGAATTTGCCCGCGAG<br>ACCTTCCAGGCCTTCCGCACCACCGACGTCGGCCGCAAGCTGATCATCGATCAGAACGTTTTTATCGAG<br>GGTACGCTGCCGATGGGTGTCGTCCGCCCGCTGACTGAAGTCGAGATGGACCATTACCGCGAGCCGTT<br>CCTGAATCCTGTTGACCGCGAGCCACTGTGGCGCTTCCCAAACGAGCTGCCAATCGCCGGTGAGCCAG<br>CGAACATCGTCGCGCTGGTCGAAGAATACATGGACTGGCTGCACCAGTCCCCTGTCCCGAAGCTGCTGT<br>TCTGGGGCACCCCAGGCGTTCTGATCCCACCGGCCGAAGCCGCTCGCCTGGCCAAAAGCCTGCCTAAC<br>TGCAAGGCTGTGGACATCGGCCCGGGTCTGAATCTGCTGCAAGAAGACAACCCGGACCTGATCGGCAG<br>CGAGATCGCGCGCTGGCTGTCTACTCTGGAGATTTCCGGCTCCGAATTCGATTACAAGGACCACGATGG<br>CGACTATAAGGATCACGACATCGACTACAAGGACGATGACGACAAGTGA |
| **LAT dCore HaloTag** | ATGGAGGAGGCCATCCTGGTCCCCTGCGTGCTGGGGCTCGCCATGTTGATGGCACTGTGTGTGCACTG<br>CCACAGACTGCCAGGATCCGGCGGCTCCGAAATCGGTACTGGCTTTCCATTCGACCCCCATTATGTGGA<br>AGTCCTGGGCGAGCGCATGCACTACGTCGATGTTGGTCCGCGCGATGGCACCCCTGTGCTGTTCCTGC<br>ACGGTAACCCGACCTCCTCCTACGTGTGGCGCAACATCATCCCGCATGTTGCACCGACCCATCGCTGCA<br>TTGCTCCAGACCTGATCGGTATGGGCAAATCCGACAAACCAGACCTGGGTTATTTCTTCGACGACCACG<br>TCCGCTTCATGGATGCCTTCATCGAAGCCCTGGGTCTGGAAGAGGTCGTCCTGGTCATTCACGACTGGG<br>GCTCCGCTCTGGGTTTCCACTGGGCCAAGCGCAATCCAGAGCGCGTCAAAGGTATTGCATTTATGGAGT<br>TCATCCGCCCTATCCCGACCTGGGACGAATGGCCAGAATTTGCCCGCGAGACCTTCCAGGCCTTCCGCA<br>CCACCGACGTCGGCCGCAAGCTGATCATCGATCAGAACGTTTTTATCGAGGGTACGCTGCCGATGGGTG<br>TCGTCCGCCCGCTGACTGAAGTCGAGATGGACCATTACCGCGAGCCGTTCCTGAATCCTGTTGACCGCG<br>AGCCACTGTGGCGCTTCCCAAACGAGCTGCCAATCGCCGGTGAGCCAGCGAACATCGTCGCGCTGGTC<br>GAAGAATACATGGACTGGCTGCACCAGTCCCCTGTCCCGAAGCTGCTGTTCTGGGGCACCCCAGGCGT<br>TCTGATCCCACCGGCCGAAGCCGCTCGCCTGGCCAAAAGCCTGCCTAACTGCAAGGCTGTGGACATCG<br>GCCCGGGTCTGAATCTGCTGCAAGAAGACAACCCGGACCTGATCGGCAGCGAGATCGCGCGCTGGCTG<br>TCTACTCTGGAGATTTCCGGCTCCGAATTCGATTACAAGGACCACGATGGCGACTATAAGGATCACGAC<br>ATCGACTACAAGGACGATGACGACAAGTGA |
| **LAT High ASA HaloTag** | ATGGAAGAGGCCATCCTGGTTCTGCTGCTGCTCCTTCTGCTTCTGCTGCCTCTTCTCTTGCTCCTCCTGC<br>TGCTGTGCGTGCACTGTCACAGATTGCCTGGATCCGGCGGCTCCGAAATCGGTACTGGCTTTCCATTCG<br>ACCCCCATTATGTGGAAGTCCTGGGCGAGCGCATGCACTACGTCGATGTTGGTCCGCGCGATGGCACC<br>CCTGTGCTGTTCCTGCACGGTAACCCGACCTCCTCCTACGTGTGGCGCAACATCATCCCGCATGTTGCA<br>CCGACCCATCGCTGCATTGCTCCAGACCTGATCGGTATGGGCAAATCCGACAAACCAGACCTGGGTTAT<br>TTCTTCGACGACCACGTCCGCTTCATGGATGCCTTCATCGAAGCCCTGGGTCTGGAAGAGGTCGTCCTG<br>GTCATTCACGACTGGGGCTCCGCTCTGGGTTTCCACTGGGCCAAGCGCAATCCAGAGCGCGTCAAAGG<br>TATTGCATTTATGGAGTTCATCCGCCCTATCCCGACCTGGGACGAATGGCCAGAATTTGCCCGCGAGAC<br>CTTCCAGGCCTTCCGCACCACCGACGTCGGCCGCAAGCTGATCATCGATCAGAACGTTTTTATCGAGGG<br>TACGCTGCCGATGGGTGTCGTCCGCCCGCTGACTGAAGTCGAGATGGACCATTACCGCGAGCCGTTCC<br>TGAATCCTGTTGACCGCGAGCCACTGTGGCGCTTCCCAAACGAGCTGCCAATCGCCGGTGAGCCAGCG<br>AACATCGTCGCGCTGGTCGAAGAATACATGGACTGGCTGCACCAGTCCCCTGTCCCGAAGCTGCTGTTC<br>TGGGGCACCCCAGGCGTTCTGATCCCACCGGCCGAAGCCGCTCGCCTGGCCAAAAGCCTGCCTAACTG<br>CAAGGCTGTGGACATCGGCCCGGGTCTGAATCTGCTGCAAGAAGACAACCCGGACCTGATCGGCAGCG |

| | |
|---|---|
| | AGATCGCGCGCTGGCTGTCTACTCTGGAGATTTCCGGCTCCGAATTCGATTACAAGGACCACGATGGCG ACTATAAGGATCACGACATCGACTACAAGGACGATGACGACAAGTGA |
| **24 Å 4TMD RFP** | ATGGGCAGCACCAGAACCGAGATCATCAGAGAGCTGGAAAGAAGCCTGCGCGAGCAGAGAGTGCTGGC CATTTTTCTGCTGGCCCTGCTGATCGTGCTGCTGTGGCTGCTGCAACAGCTGAAAGAGCTGCTGAGAGA ACTGGAACGGCTGCAGAGAGAGGGCAGCTCTGACGAAGATGTGCGGGAACTGCTGCGCGAGATCAAAG AACTGGTGGAAAACATCGTGTACCTGGTTATCATCATCATGGTGCTGGTGCTCGTGATCATTGCCCTGGC CAGAACACAGAAGTACCTGGTCGAGGAACTGAAGCGGCAGGACGGATCCGGCGGCTCCGAAATCGGTA CTGGCTTTCCATTCGACCCCCATTATGTGGAAGTCCTGGGCGAGCGCATGCACTACGTCGATGTTGGTC CGCGCGATGGCACCCCTGTGCTGTTCCTGCACGGTAACCCGACCTCCTCCTACGTGTGGCGCAACATC ATCCCGCATGTTGCACCGACCCATCGCTGCATTGCTCCAGACCTGATCGGTATGGGCAAATCCGACAAA CCAGACCTGGGTTATTTCTTCGACGACCACGTCCGCTTCATGGATGCCTTCATCGAAGCCCTGGGTCTG GAAGAGGTCGTCCTGGTCATTCACGACTGGGGCTCCGCTCTGGGTTTCCACTGGGCCAAGCGCAATCC AGAGCGCGTCAAAGGTATTGCATTTATGGAGTTCATCCGCCCTATCCCGACCTGGGACGAATGGCCAGA ATTTGCCCGCGAGACCTTCCAGGCCTTCCGCACCACCGACGTCGGCCGCAAGCTGATCATCGATCAGA ACGTTTTTATCGAGGGTACGCTGCCGATGGGTGTCGTCCGCCCGCTGACTGAAGTCGAGATGGACCATT ACCGCGAGCCGTTCCTGAATCCTGTTGACCGCGAGCCACTGTGGCGCTTCCCAAACGAGCTGCCAATC GCCGGTGAGCCAGCGAACATCGTCGCGCTGGTCGAAGAATACATGGACTGGCTGCACCAGTCCCCTGT CCCGAAGCTGCTGTTCTGGGGCACCCCAGGCGTTCTGATCCCACCGGCCGAAGCCGCTCGCCTGGCCA AAAGCCTGCCTAACTGCAAGGCTGTGGACATCGGCCCGGGTCTGAATCTGCTGCAAGAAGACAACCCG GACCTGATCGGCAGCGAGATCGCGCGCTGGCTGTCTACTCTGGAGATTTCCGGCTCCGAATTCGATTAC AAGGACCACGATGGCGACTATAAGGATCACGACATCGACTACAAGGACGATGACGACAAGTGA |
| **32 Å 4TMD RFP** | ATGGGCAGCACCAGAACCGAGATCATCACCCGGCTGAGCTTCAGCCTGCTGCTGCAACTGGTGCTGGC TATCTTTCTGCTGGCCCTGCTGATCGTGCTGCTGTGGCTGCTTCAGCAGCTGAAAGAGCTGCTGAGAGA GCTGGAACGGCTGCAGAGAGAGGGAAGCTCTGACGAGGATGTGCGGGAACTGCTGCGCGAGATCAAA GAACTGGTGGAAAACATCGTGTACCTGGTTATCATCATCATGGTGCTGGTGCTCGTGATCATTGCCCTGG CCGTGCTGCAGATGTACCTCGTCAGGGAACTGAAGCGGCAGGACGGATCCGGCGGCTCCATGGCCTCC TCCGAGGACGTCATCAAGGAGTTCATGCGCTTCAAGGTGCGCATGGAGGGCTCCGTGAACGGCCACGA GTTCGAGATCGAGGGCGAGGGCGAGGGCCGCCCCTACGAGGGCACCCAGACCGCCAAGCTGAAGGTG ACCAAGGGCGGCCCCCTGCCCTTCGCCTGGGACATCCTGTCCCCTCAGTTCCAGTACGGCTCCAAGGC CTACGTGAAGCACCCCGCCGACATCCCCGACTACTTGAAGCTGTCCTTCCCCGAGGGCTTCAAGTGGGA GCGCGTGATGAACTTCGAGGACGGCGGCGTGGTGACCGTGACCCAGGACTCCTCCCTGCAGGACGGC GAGTTCATCTACAAGGTGAAGCTGCGCGGCACCAACTTCCCCTCCGACGGCCCCGTAATGCAGAAGAA GACCATGGGCTGGGAGGCCTCCACCGAGCGGATGTACCCCGAGGACGGCGCCCTGAAGGGCGAGATC AAGATGAGGCTGAAGCTGAAGGACGGCGGCCACTACGACGCCGAGGTCAAGACCACCTACATGGCCAA GAAGCCCGTGCAGCTGCCCGGCGCCTACAAGACCGACATCAAGCTGGACATCACCTCCCACAACGAGG ACTACACCATCGTGGAACAGTACGAGCGCGCCGAGGGCCGCCACTCCACCGGCGCCTCCGGCTCCGA ATTCGATTACAAGGACCACGATGGCGACTATAAGGATCACGACATCGACTACAAGGACGATGACGACAA GTGA |
| **40 Å 4TMD RFP** | ATGGGCAGCACCAGAACCGAGATCATCACCCGGCTGAGCTTCAGCCTGCTGCTGCAACTGGTGCTGGC TATCTTTCTGCTGGCCCTGCTGATCGTGCTGCTGGTGCTCCTGATCTACCTGAAAGAGCTGCTGAGAGA GCTGGAACGGCTGCAGAGAGAGGGAAGCTCTGACGAGGATGTGCGAGAACTGCTGCGCGAGATCAAGT GGCTGGTCATCGTGATCGTGGCCCTGGTTATCATCATCATGGTGCTGGTCCTGGTCATCATTGCCCTGG CCGTGCTGCAGATGTACCTCGTGCGGGAACTGAAGAGACAGGACGGATCCGGCGGCTCCATGGCCTCC TCCGAGGACGTCATCAAGGAGTTCATGCGCTTCAAGGTGCGCATGGAGGGCTCCGTGAACGGCCACGA GTTCGAGATCGAGGGCGAGGGCGAGGGCCGCCCCTACGAGGGCACCCAGACCGCCAAGCTGAAGGTG ACCAAGGGCGGCCCCCTGCCCTTCGCCTGGGACATCCTGTCCCCTCAGTTCCAGTACGGCTCCAAGGC CTACGTGAAGCACCCCGCCGACATCCCCGACTACTTGAAGCTGTCCTTCCCCGAGGGCTTCAAGTGGGA GCGCGTGATGAACTTCGAGGACGGCGGCGTGGTGACCGTGACCCAGGACTCCTCCCTGCAGGACGGC GAGTTCATCTACAAGGTGAAGCTGCGCGGCACCAACTTCCCCTCCGACGGCCCCGTAATGCAGAAGAA GACCATGGGCTGGGAGGCCTCCACCGAGCGGATGTACCCCGAGGACGGCGCCCTGAAGGGCGAGATC AAGATGAGGCTGAAGCTGAAGGACGGCGGCCACTACGACGCCGAGGTCAAGACCACCTACATGGCCAA GAAGCCCGTGCAGCTGCCCGGCGCCTACAAGACCGACATCAAGCTGGACATCACCTCCCACAACGAGG ACTACACCATCGTGGAACAGTACGAGCGCGCCGAGGGCCGCCACTCCACCGGCGCCTCCGGCTCCGA ATTCGATTACAAGGACCACGATGGCGACTATAAGGATCACGACATCGACTACAAGGACGATGACGACAA GTGA |
| **24 Å 12TMD RFP** | ATGACCGAGAACGAGATCCGGAAGCTGAGAAAGCTGCTGCGGATCGCTATGTTCCTGCTGGTGTTTCTG CTGATCTGGACCTGGATCAGCCTGGAAACCAGCAAGACCGACGACGACCCTAGCGCTCAGTCTGAAGC TCTGGTGGCCATGAGCCTGATGCTGATTGCCGCCAGCCTGCTGATCATTGCCAAGAGCAAGCTGATGAA GTCCCGGAACGGAGGATCCGGCGGCTCCGAAATCGGTACTGGCTTTCCATTCGACCCCCATTATGTGGA AGTCCTGGGCGAGCGCATGCACTACGTCGATGTTGGTCCGCGCGATGGCACCCCTGTGCTGTTCCTGC ACGGTAACCCGACCTCCTCCTACGTGTGGCGCAACATCATCCCGCATGTTGCACCGACCCATCGCTGCA TTGCTCCAGACCTGATCGGTATGGGCAAATCCGACAAACCAGACCTGGGTTATTTCTTCGACGACCACG TCCGCTTCATGGATGCCTTCATCGAAGCCCTGGGTCTGGAAGAGGTCGTCCTGGTCATTCACGACTGGG GCTCCGCTCTGGGTTTCCACTGGGCCAAGCGCAATCCAGAGCGCGTCAAAGGTATTGCATTTATGGAGT TCATCCGCCCTATCCCGACCTGGGACGAATGGCCAGAATTTGCCCGCGAGACCTTCCAGGCCTTCCGCA CCACCGACGTCGGCCGCAAGCTGATCATCGATCAGAACGTTTTTATCGAGGGTACGCTGCCGATGGGTG TCGTCCGCCCGCTGACTGAAGTCGAGATGGACCATTACCGCGAGCCGTTCCTGAATCCTGTTGACCGCG AGCCACTGTGGCGCTTCCCAAACGAGCTGCCAATCGCCGGTGAGCCAGCGAACATCGTCGCGCTGGTC GAAGAATACATGGACTGGCTGCACCAGTCCCCTGTCCCGAAGCTGCTGTTCTGGGGCACCCCAGGCGT TCTGATCCCACCGGCCGAAGCCGCTCGCCTGGCCAAAAGCCTGCCTAACTGCAAGGCTGTGGACATCG |

| | |
|---|---|
| | GCCCGGGTCTGAATCTGCTGCAAGAAGACAACCCGGACCTGATCGGCAGCGAGATCGCGCGCTGGCTG<br>TCTACTCTGGAGATTTCCGGCTCCGAATTCGATTACAAGGACCACGATGGCGACTATAAGGATCACGAC<br>ATCGACTACAAGGACGATGACGACAAGTGA |
| **40 Å 12TMD RFP** | ATGACCAAGAAAATCATCATGGTGCTGATCCTGCTGCTGATCATTGCTATGCTGCTGCTGGTGTTCCTGC<br>TGATCGCCACCGTGGTTTCCCTGTGGTGGTCCTGGACCGACGATGATCCTAGCGCCATTTCTGAAGCCC<br>TGGTGGCCATGAGCCTGATGCTGATTGCTGCCAGCCTGCTGATTATCGCCATCAGCAAGCTGCTGAAGT<br>CCAAGAACGGCGGATCCGGCGGCTCCATGGCCTCCTCCGAGGACGTCATCAAGGAGTTCATGCGCTTC<br>AAGGTGCGCATGGAGGGCTCCGTGAACGGCCACGAGTTCGAGATCGAGGGCGAGGGCGAGGGCCGC<br>CCCTACGAGGGCACCCAGACCGCCAAGCTGAAGGTGACCAAGGGCGGCCCCCTGCCCTTCGCCTGGG<br>ACATCCTGTCCCCTCAGTTCCAGTACGGCTCCAAGGCCTACGTGAAGCACCCCGCCGACATCCCCGACT<br>ACTTGAAGCTGTCCTTCCCCGAGGGCTTCAAGTGGGAGCGCGTGATGAACTTCGAGGACGGCGGCGTG<br>GTGACCGTGACCCAGGACTCCTCCCTGCAGGACGGCGAGTTCATCTACAAGGTGAAGCTGCGCGGCAC<br>CAACTTCCCCTCCGACGGCCCCGTAATGCAGAAGAAGACCATGGGCTGGGAGGCCTCCACCGAGCGGA<br>TGTACCCCGAGGACGGCGCCCTGAAGGGCGAGATCAAGATGAGGCTGAAGCTGAAGGACGGCGGCCA<br>CTACGACGCCGAGGTCAAGACCACCTACATGGCCAAGAAGCCCGTGCAGCTGCCCGGCGCCTACAAGA<br>CCGACATCAAGCTGGACATCACCTCCCACAACGAGGACTACACCATCGTGGAACAGTACGAGCGCGCC<br>GAGGGCCGCCACTCCACCGGCGCCTCCGGCTCCGAATTCGATTACAAGGACCACGATGGCGACTATAA<br>GGATCACGACATCGACTACAAGGACGATGACGACAAGTGA |
| **Soluble HaloTag** | ATGGAATTCGGCGGCTCCGAAATCGGTACTGGCTTTCCATTCGACCCCCATTATGTGGAAGTCCTGGGC<br>GAGCGCATGCACTACGTCGATGTTGGTCCGCGCGATGGCACCCCTGTGCTGTTCCTGCACGGTAACCC<br>GACCTCCTCCTACGTGTGGCGCAACATCATCCCGCATGTTGCACCGACCCATCGCTGCATTGCTCCAGA<br>CCTGATCGGTATGGGCAAATCCGACAAACCAGACCTGGGTTATTTCTTCGACGACCACGTCCGCTTCAT<br>GGATGCCTTCATCGAAGCCCTGGGTCTGGAAGAGGTCGTCCTGGTCATTCACGACTGGGGCTCCGCTC<br>TGGGTTTCCACTGGGCCAAGCGCAATCCAGAGCGCGTCAAAGGTATTGCATTTATGGAGTTCATCCGCC<br>CTATCCCGACCTGGGACGAATGGCCAGAATTTGCCCGCGAGACCTTCCAGGCCTTCCGCACCACCGAC<br>GTCGGCCGCAAGCTGATCATCGATCAGAACGTTTTTATCGAGGGTACGCTGCCGATGGGTGTCGTCCGC<br>CCGCTGACTGAAGTCGAGATGGACCATTACCGCGAGCCGTTCCTGAATCCTGTTGACCGCGAGCCACTG<br>TGGCGCTTCCCAAACGAGCTGCCAATCGCCGGTGAGCCAGCGAACATCGTCGCGCTGGTCGAAGAATA<br>CATGGACTGGCTGCACCAGTCCCCTGTCCCGAAGCTGCTGTTCTGGGGCACCCCAGGCGTTCTGATCC<br>CACCGGCCGAAGCCGCTCGCCTGGCCAAAAGCCTGCCTAACTGCAAGGCTGTGGACATCGGCCCGGG<br>TCTGAATCTGCTGCAAGAAGACAACCCGGACCTGATCGGCAGCGAGATCGCGCGCTGGCTGTCTACTCT<br>GGAGATTTCCGGCTCCGGATCCGACTACAAGGACGATGACGACAAGTGA |
| **M HaloTag** | ATGGGCAGTTCTAAGAGCAAGCCCAAGGATCCCAGCCAGCGGCGGAACAACAACAATGGACCTGTGGC<br>CACTGAATTCGGCGGCTCCGAAATCGGTACTGGCTTTCCATTCGACCCCCATTATGTGGAAGTCCTGGG<br>CGAGCGCATGCACTACGTCGATGTTGGTCCGCGCGATGGCACCCCTGTGCTGTTCCTGCACGGTAACC<br>CGACCTCCTCCTACGTGTGGCGCAACATCATCCCGCATGTTGCACCGACCCATCGCTGCATTGCTCCAG<br>ACCTGATCGGTATGGGCAAATCCGACAAACCAGACCTGGGTTATTTCTTCGACGACCACGTCCGCTTCAT<br>GGATGCCTTCATCGAAGCCCTGGGTCTGGAAGAGGTCGTCCTGGTCATTCACGACTGGGGCTCCGCTC<br>TGGGTTTCCACTGGGCCAAGCGCAATCCAGAGCGCGTCAAAGGTATTGCATTTATGGAGTTCATCCGCC<br>CTATCCCGACCTGGGACGAATGGCCAGAATTTGCCCGCGAGACCTTCCAGGCCTTCCGCACCACCGAC<br>GTCGGCCGCAAGCTGATCATCGATCAGAACGTTTTTATCGAGGGTACGCTGCCGATGGGTGTCGTCCGC<br>CCGCTGACTGAAGTCGAGATGGACCATTACCGCGAGCCGTTCCTGAATCCTGTTGACCGCGAGCCACTG<br>TGGCGCTTCCCAAACGAGCTGCCAATCGCCGGTGAGCCAGCGAACATCGTCGCGCTGGTCGAAGAATA<br>CATGGACTGGCTGCACCAGTCCCCTGTCCCGAAGCTGCTGTTCTGGGGCACCCCAGGCGTTCTGATCC<br>CACCGGCCGAAGCCGCTCGCCTGGCCAAAAGCCTGCCTAACTGCAAGGCTGTGGACATCGGCCCGGG<br>TCTGAATCTGCTGCAAGAAGACAACCCGGACCTGATCGGCAGCGAGATCGCGCGCTGGCTGTCTACTCT<br>GGAGATTTCCGGCTCCGGATCCGACTACAAGGACGATGACGACAAGTGA |
| **PM HaloTag** | ATGGGCTGCATCAAGAGCAAGCGGAAGGACAAGGACCTGGAACTGAAGCTGCGGATCCTGCAGAGCAC<br>CGTGCCTAGAGCTAGAGATCCTCCAGTGGCCACAGAATTCGGCGGCTCCGAAATCGGTACTGGCTTTCC<br>ATTCGACCCCCATTATGTGGAAGTCCTGGGCGAGCGCATGCACTACGTCGATGTTGGTCCGCGCGATG<br>GCACCCCTGTGCTGTTCCTGCACGGTAACCCGACCTCCTCCTACGTGTGGCGCAACATCATCCCGCATG<br>TTGCACCGACCCATCGCTGCATTGCTCCAGACCTGATCGGTATGGGCAAATCCGACAAACCAGACCTGG<br>GTTATTTCTTCGACGACCACGTCCGCTTCATGGATGCCTTCATCGAAGCCCTGGGTCTGGAAGAGGTCG<br>TCCTGGTCATTCACGACTGGGGCTCCGCTCTGGGTTTCCACTGGGCCAAGCGCAATCCAGAGCGCGTC<br>AAAGGTATTGCATTTATGGAGTTCATCCGCCCTATCCCGACCTGGGACGAATGGCCAGAATTTGCCCGC<br>GAGACCTTCCAGGCCTTCCGCACCACCGACGTCGGCCGCAAGCTGATCATCGATCAGAACGTTTTTATC<br>GAGGGTACGCTGCCGATGGGTGTCGTCCGCCCGCTGACTGAAGTCGAGATGGACCATTACCGCGAGCC<br>GTTCCTGAATCCTGTTGACCGCGAGCCACTGTGGCGCTTCCCAAACGAGCTGCCAATCGCCGGTGAGC<br>CAGCGAACATCGTCGCGCTGGTCGAAGAATACATGGACTGGCTGCACCAGTCCCCTGTCCCGAAGCTG<br>CTGTTCTGGGGCACCCCAGGCGTTCTGATCCCACCGGCCGAAGCCGCTCGCCTGGCCAAAAGCCTGCC<br>TAACTGCAAGGCTGTGGACATCGGCCCGGGTCTGAATCTGCTGCAAGAAGACAACCCGGACCTGATCG<br>GCAGCGAGATCGCGCGCTGGCTGTCTACTCTGGAGATTTCCGGCTCCGGATCCGACTACAAGGACGAT<br>GACGACAAGTGA |
| **G HaloTag** | ATGGACTACAAGGACGATGACGACAAGGAATTCGGCGGCTCCGAAATCGGTACTGGCTTTCCATTCGAC<br>CCCCATTATGTGGAAGTCCTGGGCGAGCGCATGCACTACGTCGATGTTGGTCCGCGCGATGGCACCCC<br>TGTGCTGTTCCTGCACGGTAACCCGACCTCCTCCTACGTGTGGCGCAACATCATCCCGCATGTTGCACC<br>GACCCATCGCTGCATTGCTCCAGACCTGATCGGTATGGGCAAATCCGACAAACCAGACCTGGGTTATTT<br>CTTCGACGACCACGTCCGCTTCATGGATGCCTTCATCGAAGCCCTGGGTCTGGAAGAGGTCGTCCTGGT<br>CATTCACGACTGGGGCTCCGCTCTGGGTTTCCACTGGGCCAAGCGCAATCCAGAGCGCGTCAAAGGTA<br>TTGCATTTATGGAGTTCATCCGCCCTATCCCGACCTGGGACGAATGGCCAGAATTTGCCCGCGAGACCT |

| | |
|---|---|
| | TCCAGGCCTTCCGCACCACCGACGTCGGCCGCAAGCTGATCATCGATCAGAACGTTTTTATCGAGGGTA<br>CGCTGCCGATGGGTGTCGTCCGCCCGCTGACTGAAGTCGAGATGGACCATTACCGCGAGCCGTTCCTG<br>AATCCTGTTGACCGCGAGCCACTGTGGCGCTTCCCAAACGAGCTGCCAATCGCCGGTGAGCCAGCGAA<br>CATCGTCGCGCTGGTCGAAGAATACATGGACTGGCTGCACCAGTCCCCTGTCCCGAAGCTGCTGTTCTG<br>GGGCACCCCAGGCGTTCTGATCCCACCGGCCGAAGCCGCTCGCCTGGCCAAAAGCCTGCCTAACTGCA<br>AGGCTGTGGACATCGGCCCGGGTCTGAATCTGCTGCAAGAAGACAACCCGGACCTGATCGGCAGCGAG<br>ATCGCGCGCTGGCTGTCTACTCTGGAGATTTCCGGCTCCGGATCCGGTGGTAGCTTTCGTAGTGATGGC<br>AAAAAGAAAAAGAAGAAATCCAAGACCAAATGCCAGCTGCTGTGA |
| **PPF HaloTag** | ATGGACTACAAGGACGATGACGACAAGGAATTCGGCGGCTCCGAAATCGGTACTGGCTTTCCATTCGAC<br>CCCCATTATGTGGAAGTCCTGGGCGAGCGCATGCACTACGTCGATGTTGGTCCGCGCGATGGCACCCC<br>TGTGCTGTTCCTGCACGGTAACCCGACCTCCTCCTACGTGTGGCGCAACATCATCCCGCATGTTGCACC<br>GACCCATCGCTGCATTGCTCCAGACCTGATCGGTATGGGCAAATCCGACAAACCAGACCTGGGTTATTT<br>CTTCGACGACCACGTCCGCTTCATGGATGCCTTCATCGAAGCCCTGGGTCTGGAAGAGGTCGTCCTGGT<br>CATTCACGACTGGGGCTCCGCTCTGGGTTTCCACTGGGCCAAGCGCAATCCAGAGCGCGTCAAAGGTA<br>TTGCATTTATGGAGTTCATCCGCCCTATCCCGACCTGGGACGAATGGCCAGAATTTGCCCGCGAGACCT<br>TCCAGGCCTTCCGCACCACCGACGTCGGCCGCAAGCTGATCATCGATCAGAACGTTTTTATCGAGGGTA<br>CGCTGCCGATGGGTGTCGTCCGCCCGCTGACTGAAGTCGAGATGGACCATTACCGCGAGCCGTTCCTG<br>AATCCTGTTGACCGCGAGCCACTGTGGCGCTTCCCAAACGAGCTGCCAATCGCCGGTGAGCCAGCGAA<br>CATCGTCGCGCTGGTCGAAGAATACATGGACTGGCTGCACCAGTCCCCTGTCCCGAAGCTGCTGTTCTG<br>GGGCACCCCAGGCGTTCTGATCCCACCGGCCGAAGCCGCTCGCCTGGCCAAAAGCCTGCCTAACTGCA<br>AGGCTGTGGACATCGGCCCGGGTCTGAATCTGCTGCAAGAAGACAACCCGGACCTGATCGGCAGCGAG<br>ATCGCGCGCTGGCTGTCTACTCTGGAGATTTCCGGCTCCGGATCCGGTGGTAGCATGAAAAAGCATCGT<br>TGTAAATGTTGCAGCATTATGTGA |

*The WT-LAT HaloTag construct contains a 5mer glycine-serine linker between the C-terminal of the LAT region used here and the N-terminal of HaloTag. All other HaloTag constructs contain a 3mer glycine-serine linker at this location.

### Supplementary Table 6. DNA sequences for transcription factor delivery studies.

| Construct | DNA sequence |
|---|---|
| **Soluble synTF** | TGGAATTCGGCGGCTCCCCTAAGAAAAAGCGCAAAGTCTCCGGAAGCCAGTACCTGCCTGACACCGA<br>CGACCGGCACAGAATCGAGGAAAAGCGGAAGCGGACCTACGAGACATTCAAGAGCATCATGAAGAAG<br>TCCCCATTCAGCGGCCCCACCGATCCTAGACCTCCACCTAGAAGAATCGCCGTGCCTAGCAGATCCA<br>GCGCCTCTGTGCCTAAACCTGCTCCTCAGCCTTATCCTTTCACCAGCAGCCTGAGCACCATCAACTAC<br>GACGAGTTCCCCACAATGGTGTTCCCCAGCGGACAGATCAGCCAGGCTTCTGCTCTTGCTCCAGCTC<br>CTCCTCAGGTTCTGCCTCAAGCTCCTGCTCCGGCTCCAGCACCAGCTATGGTTTCTGCTTTGGCCCAG<br>GCTCCTGCACCTGTGCCTGTTCTTGCTCCTGGACCACCTCAGGCTGTTGCTCCACCAGCTCCTAAACC<br>TACACAGGCCGGCGAGGGAACACTGTCTGAAGCCCTGCTGCAACTCCAGTTCGACGACGAGGATCTG<br>GGAGCACTGCTGGGCAATAGCACAGACCCTGCCGTGTTTACCGATCTGGCCAGCGTGGACAACAGCG<br>AGTTTCAGCAGCTCCTGAACCAGGGCATCCCTGTGGCTCCTCACACCACAGAGCCCATGCTGATGGA<br>ATACCCCGAGGCCATCACCAGACTGGTCACCGGCGCTCAAAGACCTCCAGATCCTGCACCAGCACCT<br>CTTGGAGCACCTGGCCTGCCTAATGGACTGCTGAGCGGAGATGAGGACTTCAGCTCTATCGCCGACA<br>TGGATTTCAGCGCCCTGCTCGgttccGGTGGTGGCGGGTCTGGAGGAGGAGGTAGTGGTGGAGGTGGC<br>TCTggtacCGCTAGACCCGGCGAAAGACCTTTCCAGTGCCGGATCTGCATGAGGAACTTCAGCAAGGGC<br>GAGAGACTCGTGCGGCACACCAGAACACACACAGGCGAGAAGCCCTTCCAGTGTAGAATCTGTATGC<br>GCAACTTCAGCCGGATGGACAACCTGAGCACCCACCTGAGAACCCATACCGGGGAGAAGCCATTTCA<br>ATGCCGCATCTGTATGAGAAATTTTTCCCGGAAGGACGCCCTGAACCGGCACCTGAAAACACACCTGA<br>GAGGCAGCGGCTCCGGATCCGACTACAAGGACGATGACGACAAGTGA |
| **M synTF** | ATGGGCAGTTCTAAGAGCAAGCCCAAGGAcCCtAGCCAGCGGCGGAACAACAACAATGGACCTGTGG<br>CCACTGAATTCGGCGGCTCCCCTAAGAAAAAGCGCAAAGTCTCCGGAAGCCAGTACCTGCCTGACAC<br>CGACGACCGGCACAGAATCGAGGAAAAGCGGAAGCGGACCTACGAGACATTCAAGAGCATCATGAAG<br>AAGTCCCCATTCAGCGGCCCCACCGATCCTAGACCTCCACCTAGAAGAATCGCCGTGCCTAGCAGAT<br>CCAGCGCCTCTGTGCCTAAACCTGCTCCTCAGCCTTATCCTTTCACCAGCAGCCTGAGCACCATCAAC<br>TACGACGAGTTCCCCACAATGGTGTTCCCCAGCGGACAGATCAGCCAGGCTTCTGCTCTTGCTCCAG<br>CTCCTCCTCAGGTTCTGCCTCAAGCTCCTGCTCCGGCTCCAGCACCAGCTATGGTTTCTGCTTTGGCC<br>CAGGCTCCTGCACCTGTGCCTGTTCTTGCTCCTGGACCACCTCAGGCTGTTGCTCCACCAGCTCCTAA<br>ACCTACACAGGCCGGCGAGGGAACACTGTCTGAAGCCCTGCTGCAACTCCAGTTCGACGACGAGGAT<br>CTGGGAGCACTGCTGGGCAATAGCACAGACCCTGCCGTGTTTACCGATCTGGCCAGCGTGGACAACA<br>GCGAGTTTCAGCAGCTCCTGAACCAGGGCATCCCTGTGGCTCCTCACACCACAGAGCCCATGCTGAT<br>GGAATACCCCGAGGCCATCACCAGACTGGTCACCGGCGCTCAAAGACCTCCAGATCCTGCACCAGCA<br>CCTCTTGGAGCACCTGGCCTGCCTAATGGACTGCTGAGCGGAGATGAGGACTTCAGCTCTATCGCCG<br>ACATGGATTTCAGCGCCCTGCTCGgttccGGTGGTGGCGGGTCTGGAGGAGGAGGTAGTGGTGGAGGT<br>GGCTCTggtacCGCTAGACCCGGCGAAAGACCTTTCCAGTGCCGGATCTGCATGAGGAACTTCAGCAAG<br>GGCGAGAGACTCGTGCGGCACACCAGAACACACACAGGCGAGAAGCCCTTCCAGTGTAGAATCTGTA<br>TGCGCAACTTCAGCCGGATGGACAACCTGAGCACCCACCTGAGAACCCATACCGGGGAGAAGCCATT<br>TCAATGCCGCATCTGTATGAGAAATTTTTCCCGGAAGGACGCCCTGAACCGGCACCTGAAAACACACC<br>TGAGAGGCAGCGGCTCCGGATCCGACTACAAGGACGATGACGACAAGTGA |
| **PM synTF** | ATGGGCTGCATCAAGAGCAAGCGGAAGGACAAGGACCTGGAACTGAAGCTGagaATCCTGCAGAGCA<br>CCGTGCCTAGAGCTAGAGATCCTCCAGTGGCCACAGAATTCGGCGGCTCCCCTAAGAAAAAGCGCAA<br>AGTCTCCGGAAGCCAGTACCTGCCTGACACCGACGACCGGCACAGAATCGAGGAAAAGCGGAAGCG<br>GACCTACGAGACATTCAAGAGCATCATGAAGAAGTCCCCATTCAGCGGCCCCACCGATCCTAGACCTC<br>CACCTAGAAGAATCGCCGTGCCTAGCAGATCCAGCGCCTCTGTGCCTAAACCTGCTCCTCAGCCTTAT<br>CCTTTCACCAGCAGCCTGAGCACCATCAACTACGACGAGTTCCCCACAATGGTGTTCCCCAGCGGAC<br>AGATCAGCCAGGCTTCTGCTCTTGCTCCAGCTCCTCCTCAGGTTCTGCCTCAAGCTCCTGCTCCGGCT<br>CCAGCACCAGCTATGGTTTCTGCTTTGGCCCAGGCTCCTGCACCTGTGCCTGTTCTTGCTCCTGGACC<br>ACCTCAGGCTGTTGCTCCACCAGCTCCTAAACCTACACAGGCCGGCGAGGGAACACTGTCTGAAGCC<br>CTGCTGCAACTCCAGTTCGACGACGAGGATCTGGGAGCACTGCTGGGCAATAGCACAGACCCTGCCG<br>TGTTTACCGATCTGGCCAGCGTGGACAACAGCGAGTTTCAGCAGCTCCTGAACCAGGGCATCCCTGT<br>GGCTCCTCACACCACAGAGCCCATGCTGATGGAATACCCCGAGGCCATCACCAGACTGGTCACCGGC<br>GCTCAAAGACCTCCAGATCCTGCACCAGCACCTCTTGGAGCACCTGGCCTGCCTAATGGACTGCTGA<br>GCGGAGATGAGGACTTCAGCTCTATCGCCGACATGGATTTCAGCGCCCTGCTCGgttccGGTGGTGGCG<br>GGTCTGGAGGAGGAGGTAGTGGTGGAGGTGGCTCTggtacCGCTAGACCCGGCGAAAGACCTTTCCAG<br>TGCCGGATCTGCATGAGGAACTTCAGCAAGGGCGAGAGACTCGTGCGGCACACCAGAACACACACAG<br>GCGAGAAGCCCTTCCAGTGTAGAATCTGTATGCGCAACTTCAGCCGGATGGACAACCTGAGCACCCA<br>CCTGAGAACCCATACCGGGGAGAAGCCATTTCAATGCCGCATCTGTATGAGAAATTTTTCCCGGAAGG<br>ACGCCCTGAACCGGCACCTGAAAACACACCTGAGAGGCAGCGGCTCCGGATCCGACTACAAGGACG<br>ATGACGACAAGTGA |
| **G synTF** | ATGGACTACAAGGACGATGACGACAAGGAATTCGGCGGCTCCCCTAAGAAAAAGCGCAAAGTCTCCG<br>GAAGCCAGTACCTGCCTGACACCGACGACCGGCACAGAATCGAGGAAAAGCGGAAGCGGACCTACG<br>AGACATTCAAGAGCATCATGAAGAAGTCCCCATTCAGCGGCCCCACCGATCCTAGACCTCCACCTAGA<br>AGAATCGCCGTGCCTAGCAGATCCAGCGCCTCTGTGCCTAAACCTGCTCCTCAGCCTTATCCTTTCAC<br>CAGCAGCCTGAGCACCATCAACTACGACGAGTTCCCCACAATGGTGTTCCCCAGCGGACAGATCAGC<br>CAGGCTTCTGCTCTTGCTCCAGCTCCTCCTCAGGTTCTGCCTCAAGCTCCTGCTCCGGCTCCAGCACC<br>AGCTATGGTTTCTGCTTTGGCCCAGGCTCCTGCACCTGTGCCTGTTCTTGCTCCTGGACCACCTCAGG |

| | |
|---|---|
| | CTGTTGCTCCACCAGCTCCTAAACCTACACAGGCCGGCGAGGGAACACTGTCTGAAGCCCTGCTGCA<br>ACTCCAGTTCGACGACGAGGATCTGGGAGCACTGCTGGGCAATAGCACAGACCCTGCCGTGTTTACC<br>GATCTGGCCAGCGTGGACAACAGCGAGTTTCAGCAGCTCCTGAACCAGGGCATCCCTGTGGCTCCTC<br>ACACCACAGAGCCCATGCTGATGGAATACCCCGAGGCCATCACCAGACTGGTCACCGGCGCTCAAAG<br>ACCTCCAGATCCTGCACCAGCACCTCTTGGAGCACCTGGCCTGCCTAATGGACTGCTGAGCGGAGAT<br>GAGGACTTCAGCTCTATCGCCGACATGGATTTCAGCGCCCTGCTCGgttccGGTGGTGGCGGGTCTGGA<br>GGAGGAGGTAGTGGTGGAGGTGGCTCTggtacCGCTAGACCCGGCGAAAGACCTTTCCAGTGCCGGAT<br>CTGCATGAGGAACTTCAGCAAGGGCGAGAGACTCGTGCGGCACACCAGAACACACACAGGCGAGAA<br>GCCCTTCCAGTGTAGAATCTGTATGCGCAACTTCAGCCGGATGGACAACCTGAGCACCCACCTGAGA<br>ACCCATACCGGGGAGAAGCCATTTCAATGCCGCATCTGTATGAGAAATTTTTCCCGGAAGGACGCCCT<br>GAACCGGCACCTGAAAACACACCTGAGAGGCAGCGGCTCCGGATCCGGTGGTAGCTTTCGTAGTGAT<br>GGCAAAAAGAAAAAGAAGAAATCCAAGACCAAATGCCAGCTGCTGTGA |
| **PPF synTF** | ATGGACTACAAGGACGATGACGACAAGGAATTCGGCGGCTCCCCTAAGAAAAAGCGCAAAGTCTCCG<br>GAAGCCAGTACCTGCCTGACACCGACGACCGGCACAGAATCGAGGAAAAGCGGAAGCGGACCTACG<br>AGACATTCAAGAGCATCATGAAGAAGTCCCCATTCAGCGGCCCCACCGATCCTAGACCTCCACCTAGA<br>AGAATCGCCGTGCCTAGCAGATCCAGCGCCTCTGTGCCTAAACCTGCTCCTCAGCCTTATCCTTTCAC<br>CAGCAGCCTGAGCACCATCAACTACGACGAGTTCCCCACAATGGTGTTCCCCAGCGGACAGATCAGC<br>CAGGCTTCTGCTCTTGCTCCAGCTCCTCCTCAGGTTCTGCCTCAAGCTCCTGCTCCGGCTCCAGCACC<br>AGCTATGGTTTCTGCTTTGGCCCAGGCTCCTGCACCTGTGCCTGTTCTTGCTCCTGGACCACCTCAGG<br>CTGTTGCTCCACCAGCTCCTAAACCTACACAGGCCGGCGAGGGAACACTGTCTGAAGCCCTGCTGCA<br>ACTCCAGTTCGACGACGAGGATCTGGGAGCACTGCTGGGCAATAGCACAGACCCTGCCGTGTTTACC<br>GATCTGGCCAGCGTGGACAACAGCGAGTTTCAGCAGCTCCTGAACCAGGGCATCCCTGTGGCTCCTC<br>ACACCACAGAGCCCATGCTGATGGAATACCCCGAGGCCATCACCAGACTGGTCACCGGCGCTCAAAG<br>ACCTCCAGATCCTGCACCAGCACCTCTTGGAGCACCTGGCCTGCCTAATGGACTGCTGAGCGGAGAT<br>GAGGACTTCAGCTCTATCGCCGACATGGATTTCAGCGCCCTGCTCGgttccGGTGGTGGCGGGTCTGGA<br>GGAGGAGGTAGTGGTGGAGGTGGCTCTggtacCGCTAGACCCGGCGAAAGACCTTTCCAGTGCCGGAT<br>CTGCATGAGGAACTTCAGCAAGGGCGAGAGACTCGTGCGGCACACCAGAACACACACAGGCGAGAA<br>GCCCTTCCAGTGTAGAATCTGTATGCGCAACTTCAGCCGGATGGACAACCTGAGCACCCACCTGAGA<br>ACCCATACCGGGGAGAAGCCATTTCAATGCCGCATCTGTATGAGAAATTTTTCCCGGAAGGACGCCCT<br>GAACCGGCACCTGAAAACACACCTGAGAGGCAGCGGCTCCGGATCCGGTGGTAGCATGAAAAAGCAT<br>CGTTGTAAATGTTGCAGCATTATGTGA |

**Supplementary Table 7. Western blot antibodies used in this study and associated sample preparation considerations.**

| Antibody target | Supplier (#) | Denature temperature / time | Antibody dilution | Reducing/Non-reducing Laemmli | Animal of origin |
|---|---|---|---|---|---|
| FLAG tag | Sigma (F1804) | 70°C / 10 min | 1:1000 | Reducing | Mouse |
| CD9 | Santa Cruz (sc-13118) | 95°C / 10 min | 1:500 | Reducing | Mouse |
| CD81 | Santa Cruz (sc-23962) | 95°C / 10 min | 1:500 | Non-reducing | Mouse |
| Alix | Abcam (Ab117600) | 95°C / 10 min | 1:500 | Reducing | Mouse |
| Calnexin | Abcam (Ab22595) | 70°C / 10 min | 1:1000 | Reducing | Rabbit |
| Rabbit | Invitrogen (32460) | Not applicable | 1:3000 | Not applicable | Goat |
| Mouse | Cell Signaling Technology (7076) | Not applicable | 1:3000 | Not applicable | Horse |

**Supplementary Table 8. BD Fortessa flow cytometry lasers and photomultiplier tube (PMT) filter sets for specific fluorophores evaluated in this study.**

| Fluorophore | Channel name | Excitation laser (nm) | Filter set* |
|---|---|---|---|
| eBFP2 | Pacific Blue | 405 | 450/50 |
| dsRed-Express2 | PE-Texas Red | 552 | 610/20; 600LP |
| miRFP720 | Alexa 750 | 685 | 730/45;690 LP |

*LP, Long pass
